# Holding the arm still through integration of cortical commands

**DOI:** 10.1101/556282

**Authors:** Scott T. Albert, Alkis M. Hadjiosif, Jihoon Jang, Andrew Zimnik, Mark M. Churchland, John W. Krakauer, Reza Shadmehr

## Abstract

Every movement ends in a period of stillness. In current models of reaching, the commands that hold the arm still at a target, depend on the spatial location of the target, but not on the reach commands that moved the arm to the target. Contrary to this assumption, in a series of tasks in humans and non-human primates, we find that the commands that hold the arm still have a consistent mathematical relationship to the reach commands that moved the arm. With damage to the corticospinal tract, reach commands become impaired as expected, but remarkably, the mathematical dependence of hold commands upon the now imperfect reach commands remains intact. Therefore, we find that the brain uses a design principle in which its holding controller is not driven by the spatial location of the target. Rather, holding is obtained via mathematical integration of moving, potentially through a subcortical structure.

## Introduction

*Posture accompanies [movement] “like a shadow”*.

Sir Charles Sherrington

Reach commands that move the arm depend on the motor cortex^1,2^, but little is known about the postural controller^3,4^ that maintains arm position after the reach ends. Current models of reaching assume that the cortex uses its sensory representation of a target location, to generate motor commands that first move the arm, and then hold it still. This parallel architecture (Fig. 1B), where moving and holding are separately controlled, is implicitly assumed in all optimal control formulations of reaching^5–8^ and forms the basis for decoding activity of neurons in the motor cortex^9^.

**Figure 1.**
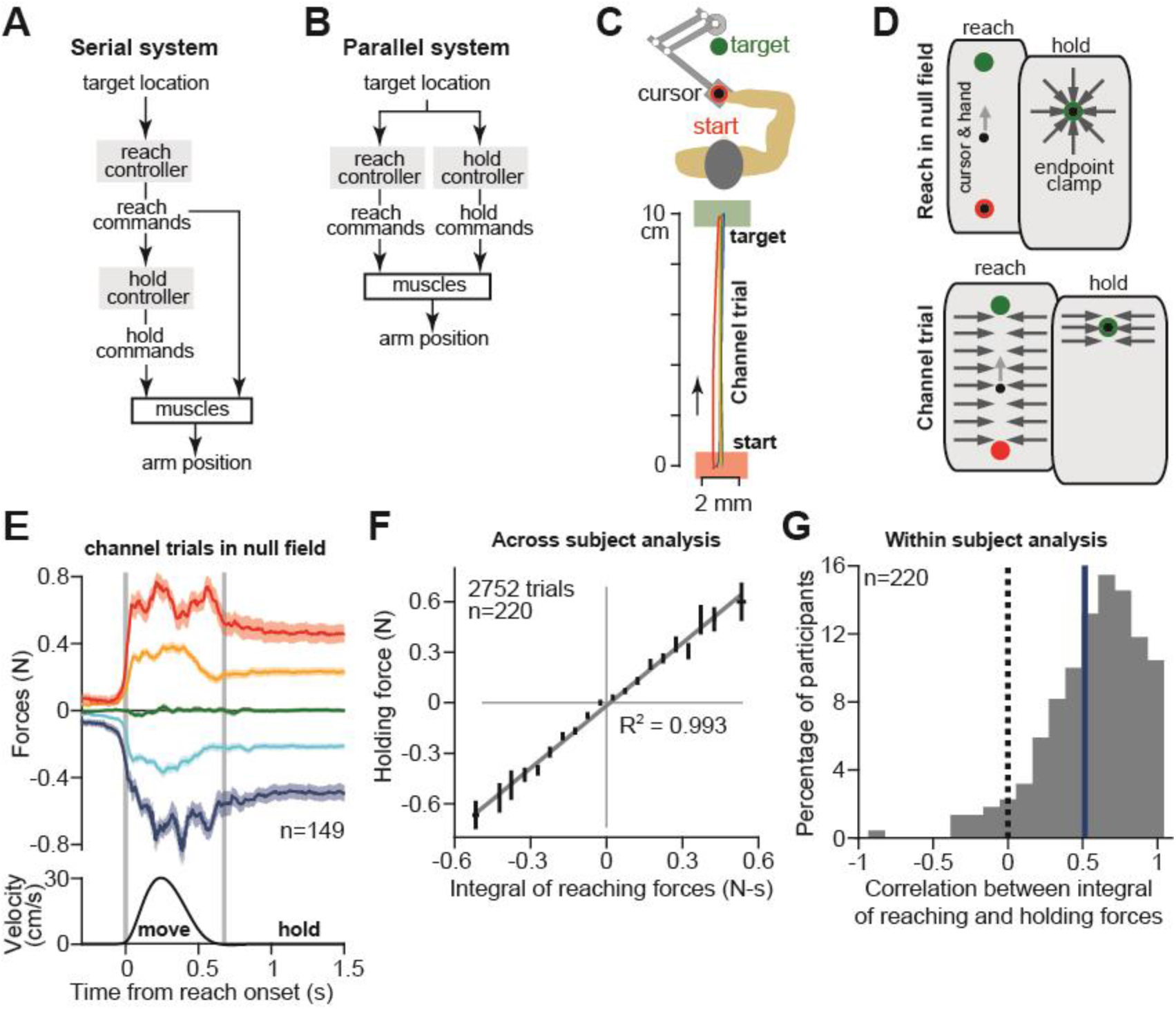
Holding commands are coupled to moving commands in human reaching. **A.** A serial architecture for the control of reaching. **B.** A parallel architecture for the control of reaching. **C.** Human participants held the handle of a robotic arm and made point-to-point reaching movements (top). On some trials, the trajectory was constrained to a straight channel (bottom) that guided the hand along virtually identical trajectories. **D.** On most trials, participants reached freely to the target. After the reach ended, an endpoint clamp held the hand in place (top). Every few trials, the channel was applied to measure lateral forces (bottom). **E.** Reaching and holding in channel trials. In the channel, natural variability in the execution of reaching forces could be measured. The mean velocity trace for all reaching movements is shown at bottom. We separated trials into five groups depending on the magnitude and sign of lateral reaching forces. Vertical gray bars indicate movement onset and termination. **F.** We calculated the integral of reaching forces, and the mean value of the holding force over a 900 ms period. We sorted trials based on the time-integral of reach forces. The line denotes a linear regression across all bins. **G.** We calculated the correlation coefficient between the time-integral of reach forces and holding forces across all channel trials in the null field, within each individual. The vertical blue line denotes the mean of the distribution. In **E** and **F**, values are mean ± SEM across all participants.

Here we present data that is inconsistent with this model. To account for the new data, we consider a different model, one that is inspired by the gaze-holding circuits of the oculomotor system.

To move the eye and then hold it steady on a target, the oculomotor system uses two separate controllers that are connected in series^10–17^ (Fig. 1A). The moving controller for a saccadic eye movement produces a set of motor commands that displace the eye^15^. Simultaneously, these commands are mathematically integrated in time by a separate brainstem structure, yielding sustained commands that hold the eye still when the saccade ends^10,14^.

Might the brain also use a serial architecture in its control of reaching, one in which the commands for holding still are not determined by the target location, but through integration of the motor commands that moved the arm?

We studied reaching and holding in both humans and non-human primates. In support of the serial model, we found that a consistent mathematical function related moving commands with subsequent holding commands, independent of the location of the target, reach duration, or direction. To test the causality of our observation, we used force fields to alter the moving commands^18^, and found a precise alteration in the commands for holding still, even though the target location never changed.

In all of our experiments, a single mathematical construct consistently captured the relationship between move commands and the subsequent hold commands; on any given trial, hold commands were proportional to the integral of the commands that moved the hand to the target. To probe the neural circuits that supported this process, we examined the movements of stroke patients that suffered from damage to the corticospinal tract (CST). Damage to the CST severely impaired move commands, but spared the integration process that led to holding.

Together, our findings suggest a new architecture for the control of reaching. While reach commands depend on the CST, holding is supported by a neural circuit outside this descending pathway. Our data support a shared design principle between movements of the arm during reaching and movements of the eye during saccades; in both the oculomotor and reach systems, moving and holding are supported by distinct neural structures that are connected to one another in series. Commands that hold the arm still depend not on the target location, but on the commands that moved the arm to its endpoint.

## Results

We designed a set of experiments to examine the relationship between commands that moved the arm and commands that held the arm still. Each of our experiments consisted of trials of point-to-point reaching, where a human or monkey moved their arm between two points in space, and then held their arm at the target location. Throughout each trial we inferred move and hold commands from the trajectory of the hand, forces the hand exerted, and finally, directly from activity of muscles in the arm. In each case, we asked if the commands for moving could predict the following commands for holding still.

### The mathematical relationship between move vs. hold commands in humans

In our human cohort, participants (n=220 in total) reached between different locations in a two-dimensional workspace while holding the handle of a robotic arm (Fig. 1C, top). On some trials, the robot produced an environment in which we could measure the lateral forces the subject produced during reaching (via a channel), as well as forces produced while holding still at the endpoint (Fig. 1D). We observed considerable trial-by-trial variability in reach forces (Fig. 1E). However, because these movements were guided within a channel, the hand followed a straight-line trajectory that always terminated at the same target location (Fig. 1C, bottom). If holding commands and moving commands were independently supported by parallel controllers (Fig. 1B) we expected that there would be no relationship between the variability present in the periods of moving, and that of holding. On the contrary, we found that on trials where subjects produced lateral reaching forces to the left or right, they also produced lateral forces in the same direction during holding (Fig. 1E). That is, even though each trial ended at the same location, variability in the holding forces was coupled to the moving forces that preceded them.

We next asked if there was a consistent mathematical function that mapped moving forces to holding forces in each of our tasks. Specifically, might the reach system employ a process of neural integration, where holding forces arise from a neural structure that integrates moving forces? To answer this question, we calculated the integral of the forces during the movement period and compared this integral to the holding forces produced after the movement ended. Remarkably, the integral of the reaching forces precisely predicted the ensuing holding forces across all participants (Fig. 1F), and also strongly predicted holding forces on a trial-by-trial basis within individuals (Fig. 1G).

### Coupling between moving and holding commands in monkeys

In our human cohort, we inferred motor commands from forces. However, a more precise measure of commands is via activity of individual muscles. Thus, we measured commands directly using intramuscular electromyography in two monkeys (Fig. 2A, left). We considered the activity of individual muscles of the arm and shoulder during point-to-point reaching in the vertical plane. On each trial, the monkey first moved its hand from a center location to one of eight targets in the vertical plane and held its hand at the target for at least 0.5 sec. Next, the monkey returned its hand back to the center location and held it there for at least 0.5 sec. Thus, from a single start point the monkey reached to one of eight targets, and then from eight different start points the monkey reached to a single endpoint.

**Figure 2.**
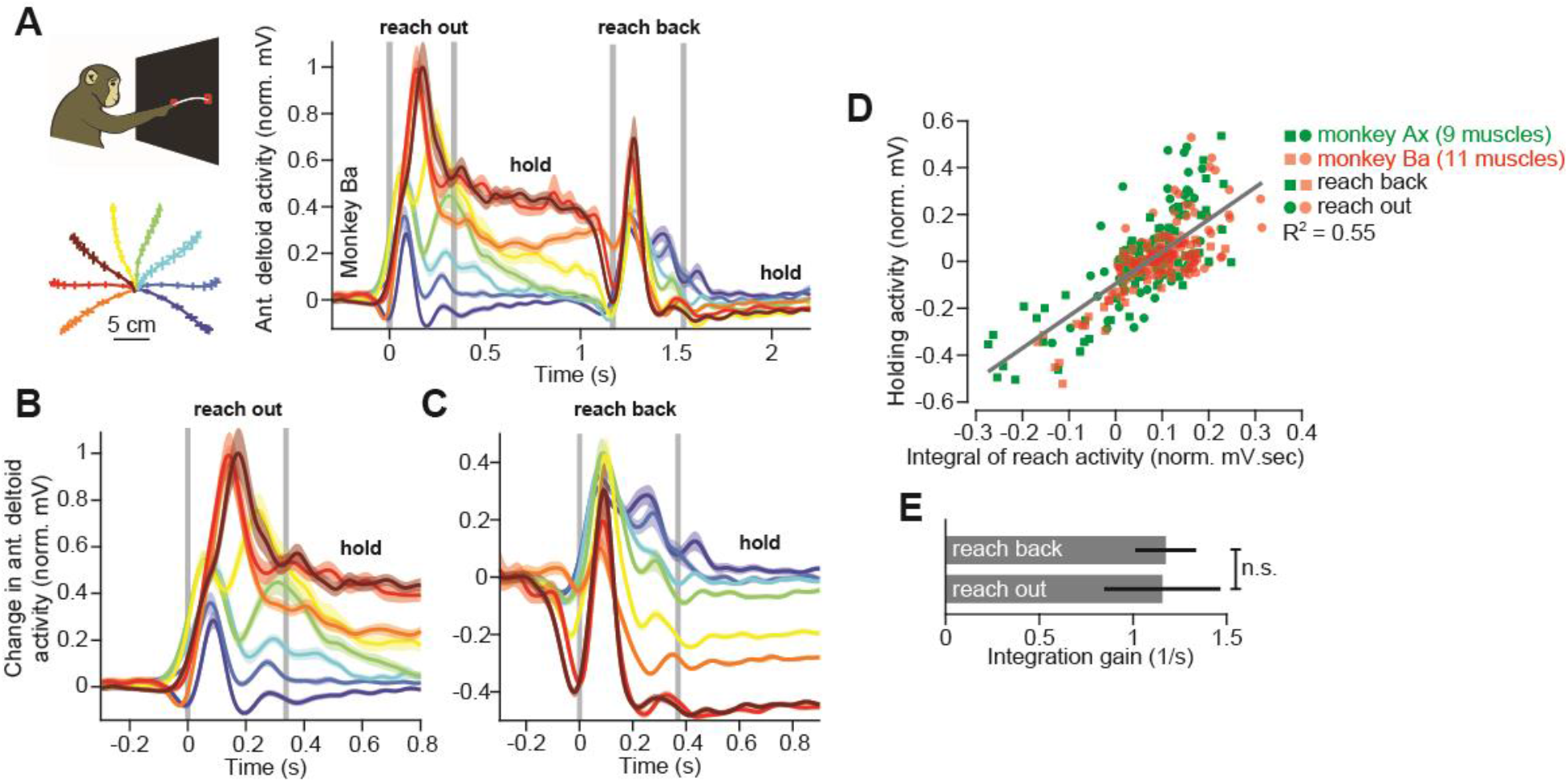
Holding commands are coupled to moving commands in non-human primate reaching. **A.** Two monkeys (top left) performed a reaching task in the vertical plane. The monkeys reached out to one of eight targets, waited, and then reached back to the home position. Trajectories for the outward reach are shown at bottom left. We measured the activity of individual muscles (n=2 monkeys, 20 muscles total) of the arm and shoulder during reaching (right). Here we show the activity of the anterior deltoid in Monkey Ba. Each color depicts reaches to a different target. **B.** We separated the activity of each muscle into two periods. The first period was the outward reach and hold. **C.** The second period was the return reach and hold. In **B** and **C**, all activity traces are referenced to the initial level of muscle activity in the holding period that preceded the reaching movement. **D.** We calculated the integral of the moving activity, as well the mean holding activity after the reach ended. We fit a linear function to the relationship between holding activity and the integral of moving activity. Data are shown for all muscles during both outward and return movements. Each point represents a single muscle and reaching movement. **E.** We separated data in **D** into the outward and return periods. We fit linear regressions to data for each individual muscle, in each period separately. Here we shown the mean slope across all muscles for each period. Statistics refer to a paired t-test. Significance was tested at the 95% confidence level. For **A, B**, and **C**, vertical gray bars indicate movement onset and termination. Values are mean ± SEM across all trials. For **E**, values are mean ± SEM across all muscles.

Each muscle showed a different pattern of activity for reaches to each of the eight targets during both moving and holding periods (Fig. 2A, right). Could holding activity be predicted by the integral of the preceding moving activity, as in our human cohort? To answer this question, we first needed to select a reference for integration. We considered the simplest hypothesis, that moving activity is integrated with respect to the sustained holding activity that preceded that movement. We separated the activity of each muscle into move and hold periods for the outwards (Fig. 2B) and backwards movements (Fig. 2C), and aligned each activity trace by subtracting pre-movement activity.

If holding activity is set by the integral of moving activity, then irrespective of start or endpoint, a muscle that increases (or decreases) activity during the reach will also have a sustained increase (or decrease) during subsequent holding. That is, on a muscle-by-muscle basis, activity during holding should be predicted from its activity during the preceding reach. Indeed, across all muscles, target locations, and reaching directions, the integral of movement period EMG was a good predictor of holding period EMG (Fig. 2D, R^2^=0.55). Remarkably, there was no difference in the integration function (i.e., the integration gain) for outward and return movements (Fig. 2E, paired t-test on single muscle regression slopes, p=0.943), nor for fast and slow movements (slow movements, 454±11ms duration; fast movements, 350±5ms; two-sample t-test on duration, p<0.001; paired t-test on integration gain, p=0.303). In summary, irrespective of reach duration, reach direction, and reach endpoint, at the level of individual muscles the mathematical integration of reach EMG predicted the ensuing hold EMG.

### Changes in moving commands lead to changes in holding commands

In both humans and monkeys, reaching commands and holding commands exhibited robust correlation, consistent with integration. To test if the relationship between moving and holding commands was causative, we designed of a set of experiments in which we externally imposed changes to the moving commands using force perturbations. If commands produced during holding directly depend on the preceding reach, changes to reach commands should alter the hold commands, even if the target location remains the same.

To test this prediction, we imposed a velocity-dependent force field during the reach (Figs. 3A and 3B), thereby encouraging adaptation that changed the reach commands (Fig. 3C, reach). Now to move straight to the target, subjects had to produce additional forces during the movement, but the period of holding still was never perturbed. Surprisingly, as reaching forces changed, so did the holding forces (Fig. 3C, hold). The holding forces were a salient feature of the holding period, persisting during the 6 second inter-trial interval (Fig. S1). Consistent with the integrator model, the holding forces of these adapted movements were accurately predicted by the integral of the preceding reaching forces (Figs. 3D and 3E).

**Figure 3.**
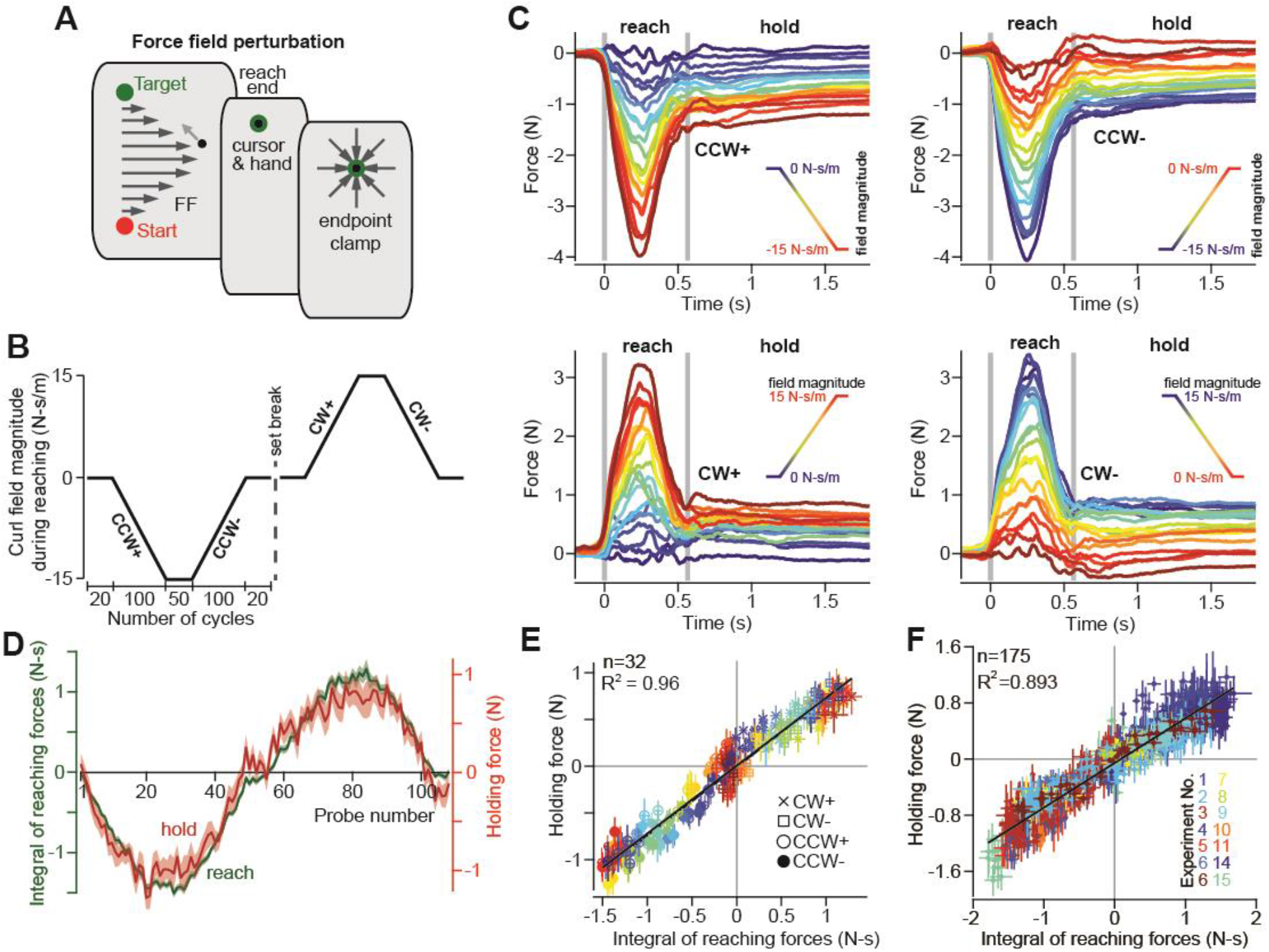
Holding forces are causally predicted by the time-integral of preceding reach forces. **A.** We exposed participants in Experiments 1, 3, and 4 (n=32) to gradually increasing (+) and decreasing (−) clockwise (CW) and counterclockwise (CCW) velocity-dependent curl force fields. **B.** The perturbation schedule. **C.** Changes in the force field magnitude caused changes in both reaching and holding forces. Each trace represents the force on one trial, averaged across participants. The vertical gray bars denote the start and end of the reaching movement. Each panel, shows consecutive trials for one of four experiment periods (CCW+, CCW−, CW+, CW−). The color of each traces indicates the force field magnitude at each point in the experiment. Blue means early in each of the four periods, and red means late. **D.** We calculated the time-integral of forces during reaching (green) and the mean holding force (red). **E.** We observed a tight coupling of holding forces to the reaching force time-integral. Each point represents one trial. Colors represent trial ordering. The shape of each marker denotes one of the four experiment periods. The solid line denotes a linear regression across all trials. **F.** We measured reaching and holding forces in 15 separate experiments. We calculated the time-integral of reaching force and holding force in each experiment. Each color denotes a different experiment. Each point represents the mean value on one trial. We fit a linear model across all trials. In **D-F**, values are mean ± SEM across participants.

To further test the causal influence of reaching on holding, we performed 14 additional experiments that employed a wide range of reaching conditions. Integration predicted holding forces when reaches started at different locations, but ended at the same target (Figs. S2A,D,E), when reaches and targets were located in various parts of the workspace (Figs. S3), when reaches were made at angles oblique to the midline of the body (Fig. S4), for reaches of larger amplitudes (Fig. S5), for forces exerted during trials where learning was generalized from other locations in the workspace (Figs. S2A,D,E), and finally in cases where moving forces alternated direction from one trial to the next (Fig. S6). Remarkably, across all of these experiments a single function accounted for approximately 90% of the variance in the data (Fig. 3F); holding forces at the endpoint were proportional to the integral of the reaching forces that brought the arm to that endpoint, irrespective of the reaching kinematics.

In a further control experiment, we considered the possibility that holding forces may be a trivial continuation of the forces applied near the end of the preceding reach, not an integration of the entire history of the reaching forces. To test this possibility, we first exposed participants (n=14) to a force perturbation that was active only during the second half of the reach (Fig. 4A, Phase 1). As expected, participants produced holding forces that increased with the integral of the moving forces (Fig. 4D, Phase 1) and remained stable across hundreds of additional trials (Fig. S5, control participants, n=11). Next, we gradually added an opposing force field during the first half of the reach (Fig. 4A, Phase 2). In this way, we guided participants to produce reaching forces that had a time-integral of approximately zero (Fig. 4B). Remarkably, as the integral of reaching forces approached zero, holding forces gradually vanished (Figs. 4D, Phase 2). That is, even though reaching forces were matched just before the end of movement (Fig. 4C, reach), the ensuing holding force differed (Fig. 4C, hold, black arrow). Reaching forces during the first half and second half of the movement made equal contributions to the final holding force; no difference in the integration function was observed between each phase of the experiment (Fig. 4E, paired t-test on slope, p=0.32, paired t-test on intercept, p=0.13). Together, these observations confirmed that holding forces were determined by integration over the entire temporal history of the preceding reach, and not simply the forces exerted near reach termination.

**Figure 4.**
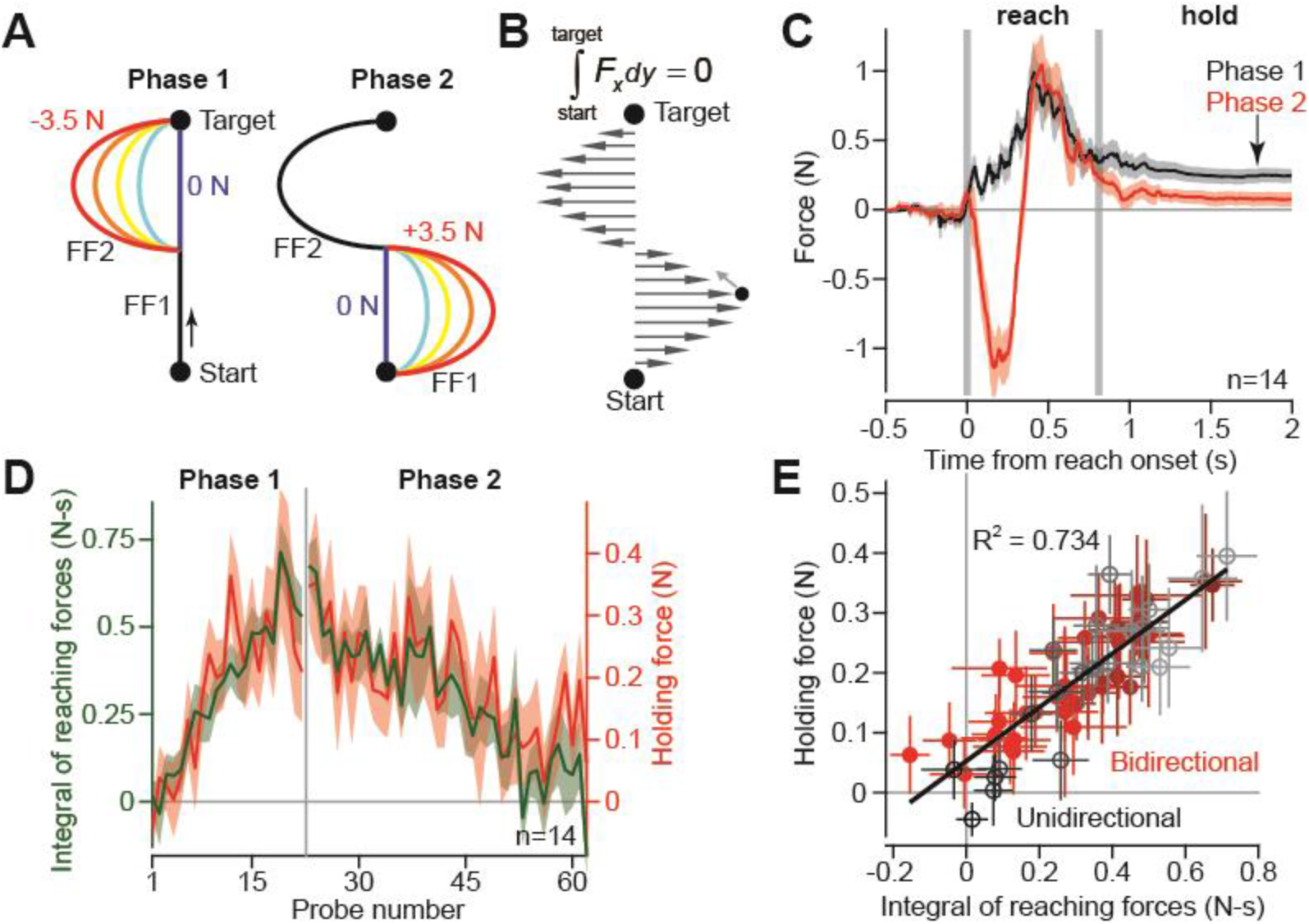
Holding forces are determined by both early and late movement commands. **A.** To test if moving forces from both the early and late parts of the reach are integrated, we performed another reaching task. Participants made 20 cm reaches between two targets. In the first phase of the experiment, a parabolic position-dependent force field was gradually applied to the second half of the reaching movement (Phase 1, left). In the second phase, an opposite force field was gradually applied to the first half of the movement (Phase 2, right). **B.** By the end of the experiment, successful compensation for the bi-directional force field yields a force time-integral of zero. **C.** Here we compare the mean force profile over several trials during Phases 1 and 2. The vertical lines denote movement onset and termination. The arrow is provided to emphasize the suppressed holding force in Phase 2. The vertical lines denote movement onset and termination. **D.** To compare reaching and holding in each phase of the experiment, we overlaid the time-integral of reaching forces (green) and holding forces (red) on each trial. The gap and vertical line denote the two separate phases of the experiment. **E.** Here we show the holding force as a function of the integral of moving force, on channel trials spaced throughout the experiment. Trial order is denoted by color (for Phase 1, black denotes early, gray denotes late; for Phase 2, darker red denotes early, and lighter red denotes late). The solid black line denotes the linear regression across all trials. In **C-E**, values are mean ± SEM across participants. In **B**, and **C**, vertical lines show movement onset and termination.

### Postural fields for the control of holding still

The experiments described thus far uncovered a heretofore unknown relationship between reaching forces and holding forces but did not explain how holding forces could be used to maintain a stable arm posture. To explore this question, participants (n=27) reached to a target as before, but now, after the reach ended, the robot slowly displaced the arm in a random direction while participants were distracted with a working memory task (Fig. 5A). In response to the displacement, the hand exerted restoring forces against the handle (Fig. 5B). When the reach took place in a null field (i.e., no perturbation to the movement), the restoring forces formed a field of postural forces (Fig. 5B, left) that had a null position near the endpoint of the preceding reach (Fig. 5E, null point of postural field). However, after participants were exposed to a force field (Fig. 5C) and changed their reach commands (Fig. 5D, reach), there was a concomitant change to the postural field (Fig. 5B, right); the null position of the postural field was no longer aligned with the target. Rather, it shifted by approximately 1 cm (Fig. 5E; paired t-test, p<10^−4^), while the orientation (Fig. 5E, paired t-test, p=0.84) and stiffness (Fig. 5E, paired t-test, p=0.62) of the field remained unchanged.

**Figure 5.**
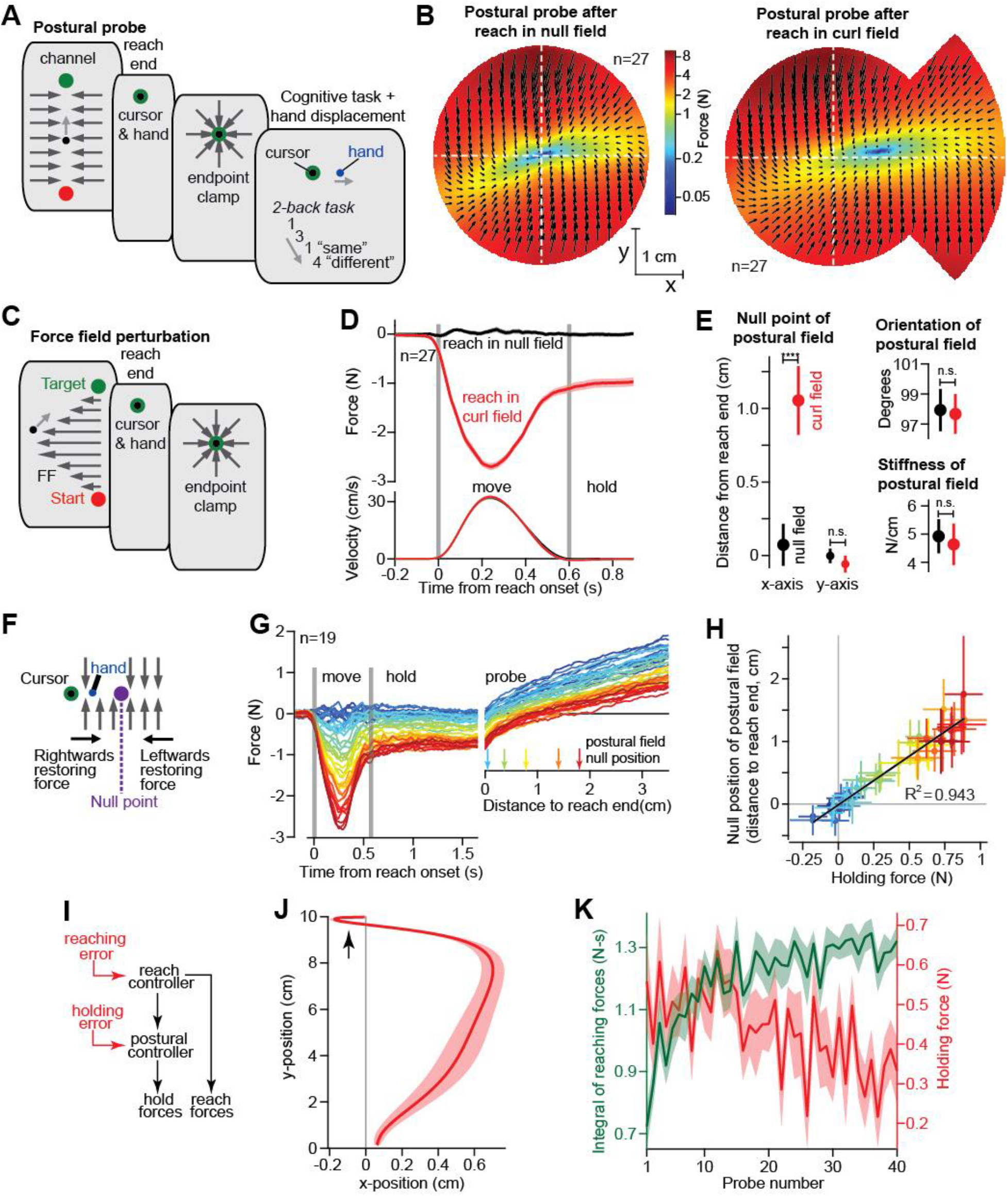
The neural integrator controls hand position through a convergent postural force field. **A.** To investigate how the neural integrator holds the arm in place we slowly displaced the hand during the holding period, while participants were distracted with a working memory task. **B.** As the hand was displaced, we measured the forces participants applied to the handle (left). We re-measured the response to holding displacements (right) after participants were exposed to a velocity-dependent curl field (force field schematic shown in C). The forces participants applied to the handle formed convergent postural force fields. Black arrows show the magnitude and direction of the restoring force for different displacements of the hand. Color reiterates the restoring force magnitude. The holding position at reach end is located at the intersection of the two dashed white lines. Forces were measured by displacing the hand outwards along 12 different lines. Interior estimates for the force were made using two-dimensional linear interpolation. **D.** We measured lateral forces (top) applied to the channel walls during reaching movements (null field period, black, and curl field period, red). **E.** We used a two-dimensional spring model to quantify postural field properties: null point, orientation, and stiffness (null field and curl field in black and red). **F.** Based on the postural fields measured in **B**, we reasoned that if the hand were moved to the null point of the postural field, restoring forces would diminish. **G.** To test this idea, participants (n=19) were exposed to a curl field that gradually increased over trials. During holding, we recorded hand forces (right) as the arm was displaced in the direction of holding forces. Arrows show the location of the null point (zero-crossing) on selected trials. **H.** We calculated the holding force before displacement of the hand, and the corresponding postural null point on each trial. Values are trial means and 95% CIs for distributions bootstrapped across participants. Linear regression was performed on the bootstrapped estimates (black line). **I.** To maintain stability at the endpoint after adaptation of the reaching controller, the postural controller must also adapt (red). **J.** In our experiments, we prevented this adaptation by removing position errors during holding with an endpoint clamp. We next performed a control experiment where this clamp was removed. In these conditions, participants experienced larger position errors while attempting to stop the hand in the target (emphasized by black arrow). **K.** The larger position errors caused adaptation in the neural integrator; now as the integral of moving forces increased, holding forces decreased. In **D, J**, and **K**, values are mean ± SEM across participants. In **D** and **G**, gray vertical bars indicate movement onset and termination. Statistics: ****p<10^−4^ and n.s. p>0.05.

This illustrated two important points. First, to hold the arm still, the nervous system created a postural force field that opposed displacement of the limb. Second, the holding forces (Fig. 5D, hold, red) we measured across all experiments in our human cohort, arose because the postural force field was misaligned with the location of the target position; the nervous system was attempting to hold their arm at a location separate from the target. If this was true, we reasoned that we could eliminate the holding force on a given trial by reducing the distance between the hand and the postural null position (Fig. 5F). Indeed, in a separate experiment, as we displaced the hand toward the postural null position, the holding force gradually approached zero, and then switched direction and grew larger as the hand was displaced beyond the postural null position (Fig. 5G). The holding force at the end of the reach scaled linearly with the distance between the hand and the postural null position (Fig. 5H). This implied that the holding force served as a proxy for the location of the null position of the postural controller; the larger the holding force, the farther the null position. Together, these experiments demonstrated that to hold the arm still, the postural controller created a position-dependent force field whose null position depended on the integral of the forces generated to move the arm to its desired endpoint.

These results create a puzzling scenario. In the presence of a force field, the reach controller readily adapts and changes the reach forces in order to steer the hand to the target. However, downstream of the reach controller, changes to the reach forces are integrated and cause the hold system to program an entirely different null position, creating a discrepancy. This implies that postural stability will be compromised in the face of an adapting reach controller. To solve this problem, the integrator must also be adaptive, possibly learning from endpoint errors that are experienced when the target is reached^19–21^ (Fig. 5I). In our experiments, the robot created an endpoint clamp (Fig. 5C) that prevented endpoint errors, thereby preventing adaptation of the hold controller. If this reasoning is correct, removal of the endpoint clamp should promote endpoint errors, thus allowing the hold controller to adapt, changing the gains of neural integration so that the null position for posture is realigned with the actual target position. This simultaneous adaptation of reaching and holding should cause holding forces at the endpoint to diminish, even as the integral of reaching forces increases.

To test this idea, we performed a control experiment in which participants (n=20) made reaching movements that ended without an endpoint clamp (Fig. 5J). As participants adapted to a force field, they now experienced larger position errors while attempting to stop the hand within the target (Fig. 5J). This increase in endpoint errors promoted adaptation in postural control; now as the integral of moving forces increased, holding forces decreased from trial to trial (Fig. 5K), in line with earlier reports^22^. We next looked for similar evidence of postural adaptation in our primary tasks. Because the endpoint clamp in our experiments was engaged only after the hand was moving sufficiently slowly, participants still committed occasional endpoint errors, albeit smaller ones. In these conditions as well, we found that individuals that made errors while attempting to slow and stop the hand at the end of the reach, ultimately produced smaller holding forces (Fig. S7). From this, we conclude that the endpoint clamp used in our primary experiments was critical for preventing adaption in the postural controller, thereby unmasking the serial link between the reach and hold systems.

### Differential contributions of the CST to reaching and holding

Our experimental results suggested that reaching and holding controllers are connected in serial through integration. One of two neural architectures could support such a network. In one case, the same cortical neurons could be responsible for generating commands and also integrating the moving commands. Alternatively, two separate populations of neurons may be responsible for movement and integration. We know that descending commands in the CST are critical for execution of voluntary movements. Does this same descending pathway also convey postural signals, or does a separate, downstream postural controller receive and then integrate the reach commands? If both reaching and holding commands are conveyed in the CST, then damage to the CST should disrupt both the generation of forces during reaching, and its integration during holding still. However, if the postural controller is downstream to the CST, then damage to CST should impair reaching, but spare the process of integration.

To examine these possibilities, we recruited stroke patients (n=13) who had suffered lesions affecting the CST pathway from the cortex through the internal capsule (Table S1). The patients exhibited profound impairments, as demonstrated by an extreme difficulty with extension of the arm during unsupported reaching (Fig. 6A, patient S015, Movie S1). With arm support (friction-less air sled), patients were better able to extend their arm at the elbow, but movements of the paretic arm continued to exhibit erratic trajectories, (Fig. 6B, patient S015; Fig. 6C, all patients), increased movement durations (Fig. 6D; paretic vs. non-paretic, paired t-test, p<0.01; paretic vs. control, two-sample t-test, p<10^−4^), and reach endpoints that terminated further away from the target location (Fig. 6E; paretic vs. non-paretic, paired t-test, p<0.01; paretic vs. control, two-sample t-test, p<0.001). The reaching impairment coincided with a marked increase in the variability of reach forces (Fig. 6F, example forces in channel trials). In fact, the time-integral of the reach forces in the paretic arm was nearly three times more variable (higher standard deviation) than healthy controls (Fig. 6G). Despite this, impaired reach trajectories still terminated in sustained holding forces (Fig. 6F, dashed lines). Remarkably, even in the most impaired movements, the within-trial coupling between the integral of reach forces and hold forces was preserved (Fig. 6H; Fig. 6I, paretic vs. non-paretic, paired t-test, p=0.776; Fig. 6I, paretic vs. control, two-sample t-test, p=0.109).

**Figure 6.**
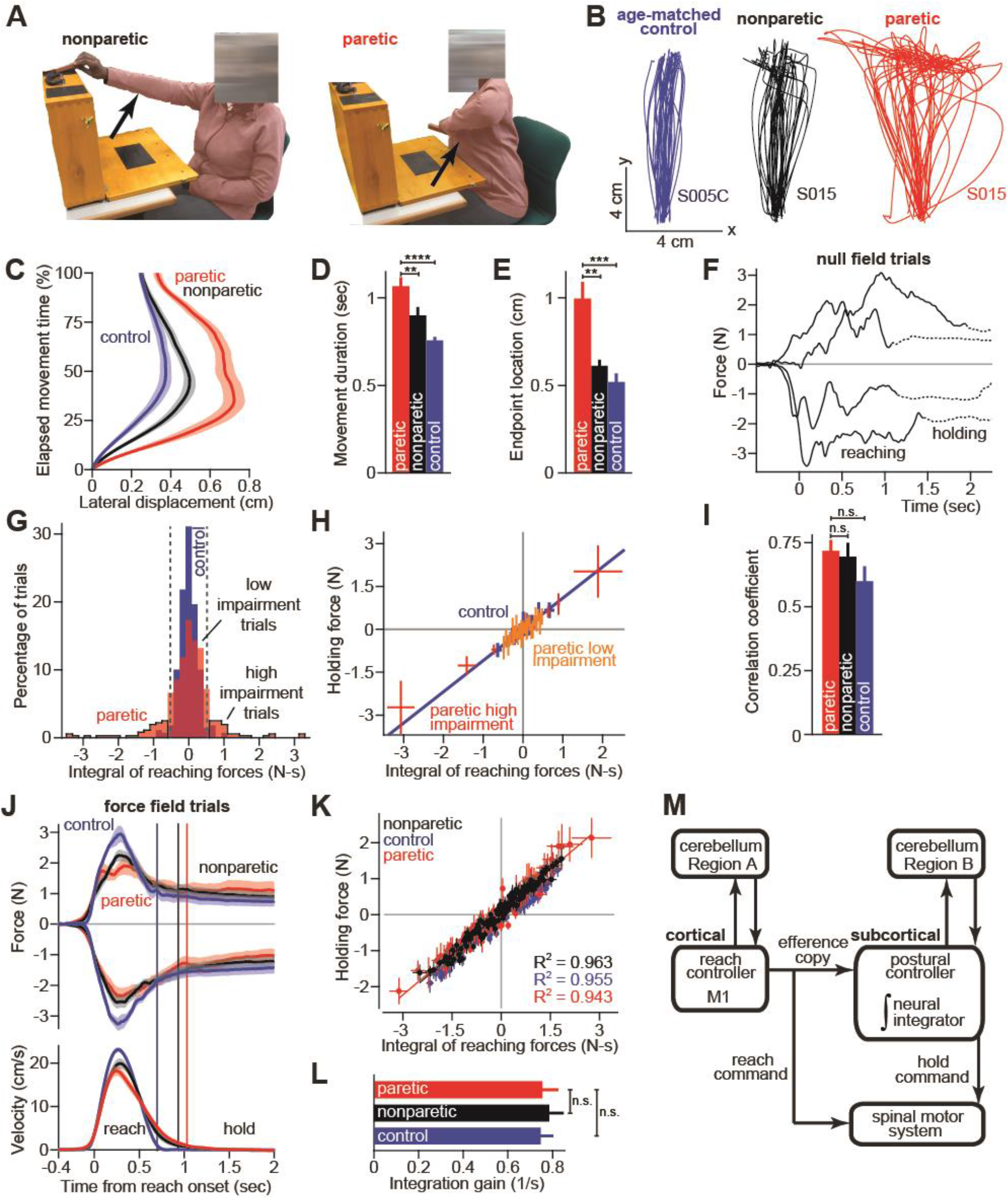
The cortical reach controller is separate from a subcortical holding integrator. **A.** Stroke survivors (n=13) participated in a set of clinical exams to measure functional impairment. Shown are isolated images for an extension-based task, for the non-paretic (top) and paretic (bottom) arms of an example participant (S015). The instruction is to place a rectangular block on the elevated surface. The task could not be completed with the paretic limb. Images show the moment of maximal extension for the paretic (bottom) and nonparetic (top) arms. **B.** To improve the range of motion of the arm, patients and healthy controls performed reaching movements holding the robotic handle, with the arm supported by an air sled. Shown are example trajectories during an initial null field period for the representative patient (black is nonparetic, red is paretic) in **A**, and a control participant (blue). **C.** We measured the distance of the hand from the desired straight-line trajectory throughout the evolution of the reach. Values are mean ± SEM across participants. **D** and **E.** We calculated several kinematic measures that captured basic features of the motor impairment in stroke: **D** shows the required duration for the reach and **E** shows the distance of the hand from the intended target at reach end. **F.** Occasionally, the arm was restricted to a straight-line path and force was measured during reaching movements in the initial null field period. Shown are example trajectories. The solid line denotes forces during moving. The dashed line denotes forces during holding still. **G.** We calculated the time-integral of moving forces during null field period trials. Shown is a histogram of moving force integrals, collapsed across all participants. To identify the most erratic reaching movements in the paretic arm (high impairment, outlined with black border), we selected reaching movements of the paretic limb that fell outside 2 standard deviations of healthy control reaches (dashed lines). **H.** We calculated the holding forces on low (orange) and high (red) impairment trials in the paretic arm, and compared these to healthy control reaches (blue). The solid blue line is a linear regression for the control participants. Lines provide estimates for the 95% confidence intervals on the mean holding force and moving force time-integral across trials binned according to the integral of moving forces. **I.** We calculated the correlation coefficient between moving force integrals and holding forces during the baseline period. Values are mean ± SEM across participants. **J.** Patients and controls were exposed to gradually increasing force fields (top is CW, bottom is CCW). Shown are the forces at the peak of the force field. Velocity traces for the hand are provided at bottom. Vertical lines show the time of movement end. Values are mean ± SEM across participants. **K.** We interrogated the relationship between the holding force and time-integral of moving force. Each point represents one trial, sorted within each subject by the time-integral of the moving force. Values are mean ± SEM across participants. **L.** We fit a linear model to capture the relationship between the moving force time-integral and holding force for each participant separately. Here we report the slope of this line, *i.e*., the gain of integration. Values are mean ± SEM across participants. **M.** A schematic of the proposed architecture for moving, holding, and neural integration. Statistics: **p<0.01, ***p<0.001, ****p<10^−4^, and n.s. p>0.05.

We next used adaptation to systematically manipulate reach forces of the patients. Because force field adaptation is largely a cerebellar-dependent process^23^, despite damage to the CST, the patients learned to alter their reach forces (Fig. 6J). Notably, we continued to observe holding forces in the paretic arm that were proportional to the integral of reach forces (Fig. 6K); remarkably, the relationship between hold forces and the integral of reach forces was identical across the paretic arm, non-paretic arm, and the dominant arm of age-matched control subjects (Fig. 6L; paretic vs. non-paretic, paired t-test p=0.758; paretic vs. control, two-sample t-test, p=0.940). Therefore, whereas damage to the CST severely affected the reach controller, it spared the integration performed by the postural controller.

## Discussion

When we make a reaching movement, a cortical system produces motor commands that move our arm towards a desired goal^24,25^, and then the reach ends in a period of holding still. But “rest is not inactivity” (Sir Charles Sherrington). In fact, to hold the arm still, muscles must receive balanced and sustained patterns of motor commands that ensure stability. Here we investigated a potential mechanism for how this stillness is achieved.

Current models assume that the sensory representation of a target provides a goal for the maintenance of posture^5–8^. However, other motor systems, like the eye^10–17^ and the head^26,27^ have found a different solution to this problem. In these systems, the commands for holding still are driven directly by the commands for movement. Here we found evidence that suggests the existence of a similar architecture that maps reaching movements into arm postures. Our experiments demonstrated that changes to reach commands, either through intrinsic sources of motor variability or adaptation, lead to subsequent changes in the commands during the hold period.

A serial model for control of reaching and holding (Fig. 6M) could account for a number of our observations. First, we discovered that holding commands, measured via activity of individual muscles or forces at the hand, had a clear mathematical relationship to moving commands, one that resembled integration. Second, variability in the moving commands, either from noise or adaptation, produced variability in the hold commands, through a process that also resembled mathematical integration. And finally, while reach commands were impaired following damage to the CST, hold commands remained coupled to move commands, through an integration process that was spared in stroke survivors.

To hold the arm at a specific location, we found that this putative reach integrator created a postural field^28,29^ that held the arm still at a location that depended not on the target location, but on the preceding reach commands. Mismatch between the center of the postural field and the actual location of the hand may help explain why reach adaptation in force fields is accompanied by paradoxical illusions in perception of arm position^30,31^.

However, holding cannot simply be an integration of moving, because moving with different dynamics (for example, different masses) should still allow one to hold at the same target position. Thus, just as the moving system can adapt to novel dynamics, the holding system should also be able to adapt to changes in the move commands. Indeed, here we found that the hold system adapted following experience of errors in the endpoint of reaching. Both reaching and holding adaptation are likely to require the cerebellum^23^, but are potentially mediated by distinct cerebellar regions^15^. The connections between the putative integrator and the cerebellum may provide a neural basis for the postural abnormalities observed in movement disorders like dystonia^32^.

Our results provide a potential solution to a critical puzzle: why does transient inhibition of the motor cortex during a reach result in “freezing of the arm” at its current posture, and not loss of muscle tone^33^? In other words, what sustains arm position against the force of gravity when the cortex is inactivated? Our proposed serial architecture for moving and holding (Figs. 1A and 6M) suggests that a separate structure, possibly located in a subcortical area, integrates the reach commands up until the moment of cortical inhibition, and then maintains posture until the reach commands resume. If this is true, stimulation of cortical areas that project to the integrator may lead to generation of specific postures, as has been found following cortical stimulation in primates^34^, and more recently in rodents^35^. However, a key experiment moving forward is to transiently stimulate the cortex during reaching and test whether this change has downstream effects on the motor commands that are generated during the subsequent period of holding still.

The idea that integration occurs outside of the CST potentially explains why decoding of neural activity from the motor cortex can provide robust predictions of reach trajectories but not arm posture during periods of holding still^9^. In addition, it is consistent with the possibility that phasic activity of neurons in the motor cortex are required for changes in arm forces, but not the sustained production of a constant force^36^. And finally, it is also in line with evidence that transient stimulation of brainstem regions in decerebrate cats, produces sustained (timescale of minutes) changes in extensor muscle force^37^.

While our findings suggest that integration occurs outside of the CST, this does not mean that cortical neurons have no contribution to the process of holding still. For example, monosynaptic projections from corticomotoneurons^38^ to alpha-motoneurons in the spinal cord are likely to be active during periods of holding still. Currently, it is unclear how this parallel pathway contributes to the maintenance of posture, and if cortical representations of posture would be generated in the cortex itself, or indirectly provided to the cortex from the putative subcortical integrator.

Understanding whether there is a subcortical system that specializes in generating commands during holding still is essential for our understanding of cortical disorders such as stroke. In fact, while integration was normal in our stroke patients, they often demonstrated abnormal arm postures at rest (Movie S1). These abnormalities could be caused by impaired cortical contributions to posture, or perhaps more provocatively, from the normal integration of chronically abnormal reach commands.

## Acknowledgements

This work was supported by grants from the National Institutes of Health (R01NS078311, R01NS095706, 1DP2NS083037, R01NS100066, 1U19NS104649, and F32NS092350), the National Science Foundation (1723967), the Simons Foundation (SCGB#542957), and the Sheikh Khalifa Stroke Institute. We would like to thank Kahori Kita for help with performing and scoring the assessments for the stroke patients, and Jennifer Keller for help performing some of these assessments and allowing us to borrow dynamometry equipment. We would also like to thank all study participants, especially stroke survivors, for their time and contribution to this work.

## Methods

Our results draw upon observations collected in 15 separate experiments. Here we first describe conceptual and experimental constructs common to each experiment, and then conclude with a description of each individual experiment.

Three populations were tested in our experiments: (1) healthy human participants, (2) stroke patients, and (3) non-human primates. Our healthy human cohort consisted of a total of n=223 individuals. Healthy participants ranged from 18-61 years of age (mean ± SD, 25.2 ± 7.9) and included 128 males and 95 females.

Our stroke patient cohort consisted of a total of n=13 older adults that had suffered damage to the corticospinal tract (CST). The stroke patients ranged from 30-80 years of age (mean ± SD, 57.8 ± 14.8) and included 6 males and 7 females. For comparison, we recruited a cohort of healthy age-matched controls who ranged from 28-81 years of age (mean ± SD, 60.6 ± 16.3) and included 5 males and 5 females. There was no significant difference in age between the patient and older healthy control populations (2-sample t-test, p=0.68). A description of lesion location and motor impairment in the stroke population is provided in our description of Experiment 15. All experiments were approved by the Institutional Review Board at the Johns Hopkins School of Medicine.

Finally, our non-human primate dataset consisted of intramuscular electromyography (EMG) and reach kinematics recorded from two monkeys (Ax and Ba) as described in^39,40^. Those data were reanalyzed here.

### Apparatus

In the experiments involving humans, participants held the handle of a planar robotic arm (Fig. 1C, top) and made point-to-point reaching movements between different target locations in the horizontal plane. The forearm was obscured from view by an opaque screen. An overhead projector displayed a small white cursor (diameter = 3mm) on the screen that tracked the motion of the hand. Visual feedback of the cursor was provided continuously throughout the entirety of each testing period, except where otherwise noted. Throughout testing we recorded the position of the robot handle using a differential encoder with submillimeter precision. We also recorded the forces produced on the handle by the subject using a 6-axis force transducer. Data were recorded at 200 Hz. Except where otherwise noted, kinematic timeseries were aligned to the onset of movement at the time point where hand velocity crossed a threshold of 1 cm/s.

### Trial structure

At trial onset, a circular target (diameter = 1 cm) appeared in the workspace, coincident with a tone that cued subject movement. Participants then voluntarily reached from the starting position to the target. After stopping the hand within the target, a holding period of various durations (1.8 to 6.5 seconds) ensued where subjects were instructed to continue holding the handle within the target. After this holding period, a random inter-trial-interval sampled uniformly between 0.3 and 0.4 seconds elapsed prior to the start of the next trial.

At the end of each reach, coincident with the start of the holding period, movement timing feedback was provided. If the preceding reach was too slow, the target turned blue and a low tone was played. If the reach was too fast, the target turned red and a low tone was played. If the reach fell within the desired movement interval (450-550ms except where otherwise noted) the target “exploded” in rings of concentric circles, a pleasing tone was played, and a point was added to a score displayed in the upper-left-hand corner of the workspace. Participants were instructed to obtain as many points as possible throughout the experimental session. In all experiments, each trial was a distinct point-to-point reaching movement, followed by a hold period.

### The reaching period

Experiments began with a set of null field trials (no perturbations from the robot). After this period, participants were exposed to force field perturbations during reaching. For the majority of experiments, the force field was a velocity-dependent curl field in which the robot generated forces proportional and perpendicular to the velocity of the hand according to:

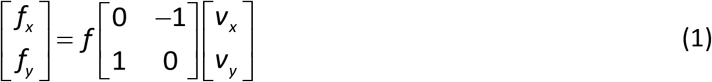

where *v_x_* and *v_y_* represent the x and y velocity of the hand, *f_x_* and *f_y_* represent the x and y force generated by the robot on the handle, and *f* represents the magnitude of the force field. We varied the force field magnitude across trials within a task. Participants were exposed to both clockwise (CW, *f* > 0) and counterclockwise (CCW, *f* < 0) curl fields. In most experiments, the perturbation was introduced in a gradual manner where the magnitude of the force field was increased very slowly from one trial to the next. In some experiments, participants were exposed to an abrupt force field where the force field magnitude immediately transitioned from zero to the maximal field strength at the start of the perturbation block.

In Experiments 8 and 9, a position-dependent force field was applied to the hand, as opposed to the curl field described in Eq. (1). We constructed the position-dependent fields from two individual force fields that were applied separately during the first (called FF_1_) and second (called FF_2_) halves of the reaching movement, as a function of displacement. Each force field was programmed as a quadratic function of distance, and applied a force solely along the horizontal axis perpendicular to the vertical reaching movement. The combined output of FF_1_ and FF_2_ produced a force field with zero force at the start position, target position, and midpoint of the reach. FF_1_ applied peak force at 25% of the displacement from start to target. FF_2_ applied peak force at 75% of the displacement from start to target (see Figs. 4A and 4B for a visual depiction). Mathematically, the entire position-dependent field can be represented as:

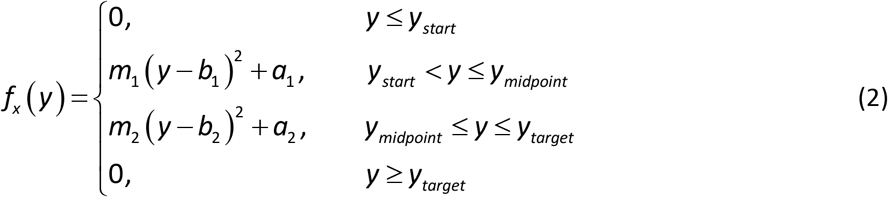

where *f_x_* represents the force applied by the robot along the axis perpendicular to the movement (the x-axis of the workspace), *y* represents the y-position of the hand, *y_start_* represents the y-coordinate of the starting position, *y_midpoint_* represents the y-coordinate of the midpoint between the starting position and the target, and *y_target_* represents the y-position of the target. The terms *m_1_, m_2_, b_1_, b_2_, a_1_*, and *a_2_* are constants that provide the appropriate shape of the position-dependent field according to:

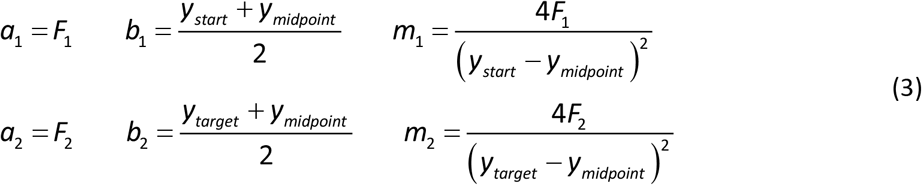

where *F_1_* and *F_2_* specify the maximum force produced during FF_1_ and FF_2_ respectively. The sign of each parameter also determines the direction of the applied force (if *F_1_* and *F_2_* are of opposite sign, the position-dependent force field applies a force to the left during one half of the reach and a force to the right during the other half of the reach). Trial-by-trial changes in *F_1_* and *F_2_* are discussed in our description of Experiments 8 and 9.

### The holding period

At the end of each reaching movement, a holding period elapsed where the hand was maintained within the target location for some period of time. Participants were instructed to simply hold the handle of the robot and wait for the start of the next trial. In most experiments, each holding period lasted for a period of at least 1.8 seconds (a fixed 1.5 seconds with an additional inter-trial-interval of at least 0.3 seconds), but sometimes lasted up to 6.5 seconds. In Experiments 5, 10, and 11, the holding period lasted only 0.5 seconds, but was followed by a long cognitive task.

In contrast to typical force field paradigms, these long inter-trial holding periods were necessary in order to test whether the integration of moving forces predicted behavior during the maintenance of posture. To isolate the serial effect of moving on holding, we attempted to remove any disturbances that could lead to adaptation of the holding system. We hypothesized that spatial errors in the stabilization and maintenance of the final hand position might provide such a teaching signal. To minimize this potential source of holding error, we applied a two-dimensional endpoint clamp. This endpoint clamp prevented motion of the hand during the hold period, despite any forces the participant might have applied to the handle. The endpoint clamp was programmed as a “well” within the target location that attracted the hand in two dimensions, with stiff spring-like mechanics (stiffness = 4000 N/m, viscosity = 75 N-s/m). The endpoint clamp was applied when the hand entered the target location and the hand velocity fell below a threshold value of 3.5 cm/s, coincident with audiovisual feedback concerning the accuracy and speed of the terminated reaching movement.

### Neural integration in human reaching movements

In our primary experiments, we asked if a process of neural integration could explain the relationship between moving and holding forces in the arm. To test this idea, we measured moving and holding forces during normal reaching movements and also during reaching movements that were adapted to a force field. For each movement, we calculated the time-integral of the forces that moved the arm during the reach, and compared this integral to the holding forces the participant applied to the handle during the holding period.

To calculate the reaching force integral, we numerically integrated the forces applied to the handle during movement. Movement onset and offset were detected by identifying the time points at which the hand crossed a velocity threshold of 1 cm/s. To calculate the holding force at the end of the reach, we computed the average force applied to the handle over a 900 ms period after the termination of the reaching movement (100-1000 ms after reach termination). To study the relationship between holding forces and the reaching force time-integral, we linearly regressed holding forces onto the moving force integral. At times, we performed this regression within single subject datasets, and at others, on quantities that were averaged across subjects. To measure the strength of reaching and holding correlations within each subject, we also calculated correlation coefficients.

Reaching and holding forces were measured on designated “channel” trials where the motion of the handle was restricted to a linear path connecting the start and target locations (Fig. 1D). To restrict hand motion to the straight-line channel trajectory, the robot applied perpendicular stiff spring-like forces with damping (stiffness = 6000 N/m, viscosity = 250 N-s/m). This channel condition was also used to measure holding forces. That is, on channel trials, the robot did not apply a two-dimensional endpoint clamp as we described earlier (see *Holding period*). Rather, the channel condition remained unchanged during holding so that the robot only maintained equilibrium of the hand along the perpendicular axis.

Reaching and holding forces were measured at regular intervals throughout the experiment. In general, every 5^th^ outwards reaching trial was performed in a channel to probe moving and holding forces. All backwards reaching movements (the reaching direction that never experienced a perturbation) were performed in either a channel trial or a partial channel trial where the channel was inactivated partway through the reach, after the hand had traveled 40% of the desired movement amplitude. By beginning each backwards reaching movement in the channel condition, we prevented the possibility that holding forces during the preceding hold period might move the hand off the target if the channel was disengaged.

Offline we isolated forces produced against the channel wall, perpendicular to the direction of the primary movement. To do this, we calculated the average force during channel trials interspersed throughout the baseline period, prior to the introduction of force perturbations. We then subtracted this baseline force time series from all of the force time series recorded during channel trials throughout the experiment. All force profiles and measurements reported herein are derived from force time series where the baseline reaching force is removed. The only exceptions to this are the forces reported during the measurement of the postural restoring field in Experiments 10 and 11.

To test if the reaching and holding systems of the arm could be linked by a neural integrator, we first considered normal reaching movements, collected during the initial period of each experiment before the introduction of any force field (Figs. 1A-D). On these trials, we found that holding forces were coupled to the reaching forces that preceded them. To test if the forces were related through a process of integration, we first subtracted the mean channel force profile as described above. What remained were residual forces relative to the mean. We calculated the integral of these residual forces during reaching, and the average residual force during holding as described above. We then sorted these residuals based on the time-integral of reaching forces. For each subject, we placed reaching force time-integrals and holding forces into one of twenty bins spanning reaching force time-integrals of −0.5 to 0.5 N-s. We then calculated the average reaching force time-integral and holding force in each bin and performed a linear regression (Fig. 1F). For visualization of these residuals, we performed the same process, but using only 5 bins (Fig. 1E). To report the strength of the moving and holding force relationship within single subjects, we calculated the correlation coefficient between these quantities for each subject (Fig. 1G).

### Neural integration in non-human primate movements

Monkeys participated in a delayed reaching task described elsewhere^39,40^. Briefly, at the start of each trial the monkey held his hand at a central home location. After a “GO” cue, the monkey executed a reaching movement to one of eight peripheral targets displayed in the vertical plane on a monitor (Fig. 2A). The monkey held his arm at the target for at least 0.5 sec. After this holding period, the monkey then returned his arm to the central home location and held the arm at the hold location for at least 0.5 sec, until the start of the next trial. On each trial, hand position was recorded through infrared optical tracking of a bead fixed to the third and fourth fingers. The activity of several muscles was also recorded using intramuscular electrodes. In Monkey Ax, these muscles included the anterior, medial, and posterior deltoid, the medial and lateral bicep, and the upper and lower trapezius. From Monkey Ba, these muscles included the anterior, medial, and posterior deltoid, the pectoralis, the brachialis, the medial and outer bicep, and the upper and lower trapezius. Prior to analysis, EMG signals were bandpass filtered (10-500 Hz), digitized at 1kHz, rectified, and finally smoothed with a Gaussian kernel (standard deviation of 20 ms). Muscle activity was then averaged across reaching movements towards each target, separately.

We asked if holding activity in each muscle could be predicted from the integral of the preceding moving activity. To ask this question, we needed to select a reference point for integration. Here we hypothesized that the natural reference point for integration is the static pre-movement activity present in the muscle during the holding period preceding movement onset. Under this hypothesis, the reference point about which moving activity is integrated is not fixed, but changes with the level of postural activity in each muscle. For each muscle we subtracted the level of activity recorded as the monkey held the hand at the home location, prior to the start of movement (from −700ms to −350ms relative to movement onset). After this subtraction, we then normalized muscles to their maximal absolute activity across movements towards each of the eight targets. This normalization process ensured that each activity traces had a maximal absolute activity of one.

We spilt each trial into two periods. The first period consisted of the outwards reaching movement and the ensuing peripheral holding period. The second period consisted of the return reaching movement and the ensuing central holding period. For the second period, to adjust the reference level for integration, we subtracted the mean holding activity that preceded the return reaching movement (−300ms to −200ms relative to movement onset of the return reaching movement) while the monkey held his hand at the peripheral target.

After separately isolating the outward and return periods (Figs. 2B and 2C), we asked if the integral of moving activity predicted holding activity, in each muscle. To calculate the integral of moving activity, we integrated normalized muscle activity across a time window of −140ms relative to movement onset, up until movement termination. We started to integrate muscle activity prior to movement onset to capture changes in muscle activity that preceded the evolution of reach kinematics. To calculate holding activity, we calculated the mean activity in each muscle over a 150ms time window beginning 300ms after the termination of movement. Of the 320 different movement types (20 muscles x 2 reaches per trial x 8 targets), 6 movements (1.9% of trials) had reach durations that were too slow to gain an accurate measurement of holding activity prior to the start of the next movement. To identify these trials, we used a cutoff for movement duration of 850ms. The 6 trials with movement durations that exceeded 850ms were not included in our analysis.

After calculating moving and holding activity, we used linear regression to determine how well the integral of moving activity predicted holding activity, across both outward and return movement periods, all muscles, and all eight targets (Fig. 2D). To determine if the integration gain differed for outward and return movements, within each individual muscle, we linearly regressed holding activity onto the integral of moving activity for the outward and return periods separately. These regressions yielded two integration gains (*i.e*. regression slope) for each muscle: one for the outward movement and one for the return. To test for a difference in the integration gains, we performed a paired t-test where each muscle served as an observation and the movement period served as the variable.

To determine if the integration gain differed for movements of different speeds, we employed a slightly different process. For each type of reaching movement (16 total, 8 targets x 2 movements per trial), we sorted muscle activity according to the movement duration. Because the activity of each muscle was not recorded simultaneously, there was natural variability across muscle recordings in the duration of the reaching movement. We separated muscle activity into two groups, a fast movement group and a slow movement group, based on the duration of the movement relative to the median movement duration. We then linearly regressed holding activity onto the integral of moving activity across the activity of each muscle in each the fast and slow groups, separately. This process yielded 16 different regression gains (one for each type of reaching movement) for both the slow and fast movement groups. To test for a difference in the integration gains, we performed a paired t-test where each movement type served as an observation and the movement speed served as the variable. In the main text, we also report the mean reach duration of all movements in the fast and slow bins. We used a two-sample t-test to test for a difference in the mean reach duration.

### Measuring the postural field

Traditionally, the holding controller is thought of as an impedance controller that is set to maintain an effector at a specific location^3,4,41^. In a subset of our experiments, we set out to characterize this impedance controller over small displacements of the hand, and understand how it might be impacted by the preceding reaching movement. Our question was, does the process of holding still at a given location depend on the motor commands that moved the arm to that location? To answer this question, we measured how hand position was controlled before and after adaptation to a force field. The force field was a tool that we used to induce participants to systematically vary their reaching force between the same two points in space.

To interrogate the holding controller, in Experiments 10 and 11, we measured how the arm reacted to displacements in its final position. To do this, we used the robot to move the hand slowly in a random direction after the hand stopped within the target. As the hand was moved, visual feedback of hand position was prevented: the display cursor was frozen at the holding location throughout the robotic intervention. To quantify the output of the holding controller, we measured the forces the subject applied to the handle while the robot moved the hand. To prevent participants from voluntarily opposing the imposed hand displacement, we distracted each participant with a difficult working memory task. We did not inform participants as to the nature or presence of the postural perturbation. Instead, we instructed participants to solely concentrate on the working memory task and obtain as many points as possible by answering memory questions correctly. Points for correct responses were combined with the points awarded for successful reaching movements.

The postural perturbation consisted of a straight-line displacement designed to make the probe as imperceptible as possible. To move the hand, we placed the hand in a two-dimensional clamp with stiff spring-like mechanics (stiffness = 4000 N/m, viscosity = 75 N-s/m) and moved the equilibrium position of the clamp through the workspace a total of either 2.5 cm, 4 cm or 5 cm, depending on the trial. The imposed motion consisted of three phases. In the first phase the hand was moved from its resting position to some desired position and velocity in accordance with a minimum jerk trajectory. In the second phase, the hand was maintained at its current velocity for some additional displacement. Finally, the hand was brought to rest over some desired displacement in accordance with a minimum jerk trajectory.

At the start of each displacement, the clamp equilibrium position was first moved a short distance (0.15 cm for 2.5 cm probes, 0.15 cm for 4 cm probes, and 0.3 cm for 5 cm probe) along a minimum jerk trajectory, over a short duration (0.75 seconds for 2.5 cm probes, 0.75 seconds for 4 cm probes, and 1.5 seconds for 5 cm probes). At the end of this displacement the velocity of the hand was equal to 0.375 cm/s. The hand was then moved at this constant velocity for a specified displacement (2.2 cm for 2.5 cm probes, 3.7 cm for 4 cm probes, and 4.4 cm for 5 cm probes). This constant velocity displacement lasted for 5.87 seconds for 2.5 cm probes, 9.87 seconds for 4 cm probes, and 11.73 seconds for 5 cm probes. After this constant velocity period, the hand was slowed to rest over a short distance (0.15 cm for 2.5 cm probes, 0.15 cm for 4 cm probes, and 0.3 cm for 5 cm probe) along a minimum jerk trajectory, over a short duration (0.75 seconds for 2.5 cm probes, 0.75 seconds for 4 cm probes, and 1.5 seconds for 5 cm probes). Finally, an additional buffer period of 0.3 seconds was added after reaching the final displaced position, prior to the end of the probe trial. The total duration of the probe was therefore 7.67 seconds for 2.5 cm probes, 11.67 seconds for 4 cm probes, and 15.03 seconds for 5 cm probes.

Critically, as stated earlier, the participant was not provided position feedback during the postural probe. Instead, cursor feedback of hand position was frozen at its holding location. Therefore, at the end of the postural probe, there was a discrepancy between the location of the hand and the location of cursor feedback. To seamlessly reunite the hand with its cursor feedback without drawing the attention of the participant, we manipulated visual feedback during the following reaching trial; as the next reach was executed, we projected the cursor position onto the line connecting the frozen cursor position and the position of the next target. In this way, it appeared to the participant as if they were reaching perfectly straight between the start and target position. At the same time, we confined the motion of the hand to a straight line connecting its displaced position with that of the next target. When the hand entered the target, a small and brief force pulse was applied to move the hand to the center of the target at which point *x* and *y* feedback was reunited with the true hand position.

### Quantifying the null point and shape of the postural field

As the hand of the participant was moved by the robot during postural probe trials, the displacement of the hand was opposed by postural restoring forces^3,4^. We used these forces to quantify how postural control of arm position responded to changes in the movement forces that transported the arm. To mathematically characterize the two-dimensional field of restoring forces, we fit a simple mathematical model^3^ that treated the arm as a linear two-dimensional spring with a single equilibrium point:

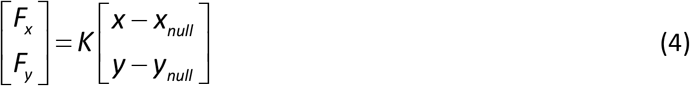

where *F_x_* and *F_y_* are the forces applied to the handle due to displacement of the hand from the null point of the system (*x_null_,y_null_*) to some position (*x,y*). The stiffness matrix *K*,

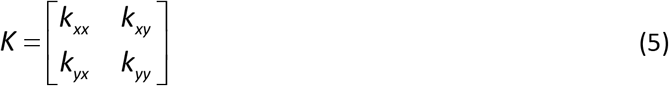

describes the magnitude and orientation of the stiffness field. We constrained *K* to be a symmetric matrix (*i.e., k_xy_ = k_yx_*). We fit this linear spring model to the postural restoring forces by identifying the parameter set (5 free parameters, *x_null_, y_null_, k_xx_, k_xy_, k_yy_*) that minimized the sum of squared error between the hand forces (collapsed across the x and y axes) predicted by Eq. (4) and the hand forces measured during all of the postural probe trials. For this fit, we used the forces measured within the ellipse bounded by −2.25 to 2.25 cm along the x-axis and −1.5 to 1.5 cm along the y-axis, relative to the final hand position. To locate the optimal parameter set, we used the genetic algorithm in MATLAB R2018a. We repeated the genetic algorithm search 8 times and selected the one that minimized the squared error cost function. The optimal parameter set provided a good fit to the data, accounting for approximately 70% of the variance in the observed postural field (R^2^ during baseline period, mean ± SEM: 0.70±0.03; R^2^ after adaptation: 0.69±0.02).

After fitting this linear model to the postural restoring field, we evaluated if the location and shape of the field changed after the reaching forces changed due to exposure to a force field. First, we looked for within-subject changes to the location of the null point (Fig. 5E, *x_null_* and *y_null_*) of the field. Second, we looked for within-subject changes to the orientation (Fig. 5E, orientation) of the field, and the overall stiffness magnitude of the field (Fig. 5E, stiffness). To calculate the orientation of the field, we considered the eigenvector of the stiffness matrix *K* corresponding to the largest eigenvalue of *K*. We calculated the angle of this eigenvector in the *x-y* plane. To compute the stiffness of the field, we calculated the Frobenius norm of the stiffness matrix *K*.

### Working memory task

In Experiments 5, 10, and 11 we employed a cognitive task to distract participants during measurement of holding forces. The working memory task consisted of a modified 2-back task where subjects were randomly shown an integer between 1 and 4. The integers appeared one at a time so that the next integer replaced the previous integer on the screen (Fig. 5A). Participants were told to determine if the integer on the display matched the integer shown two numbers in the past. If the integers matched, participants verbally responded with the keyword “same”. If the integers did not match, participants verbally responded with the keyword “different”. If the response was correct a pleasant tone was played and a point was added to the experiment score. If the response was incorrect a low pitch tone was played and no point was awarded. To confirm that participants were engaged in the cognitive task, we recorded each correct and incorrect response. Participants were clearly engaged in the cognitive task and responded correctly to 91.8 ± 0.6% of items correctly, at rates of approximately 0.77 ± 0.6 items per second.

### Measuring errors in the process of holding still

Our primary analysis demonstrated how changes to reaching forces led to changes in the process of holding still. In most cases, we induced changes in the reaching forces via imposition of a force field. We attempted to eliminate adaptation in the holding controller by applying an endpoint clamp during the holding period. These endpoint clamps, however, could not eliminate errors individuals made as they attempted to stop the hand within the target before the endpoint clamp was applied. We considered the possibility that the holding controller could be modified by these stabilization errors.

To isolate errors near the end of the reaching movement, we spatially aligned each trajectory by subtracting the final x and y coordinates of the hand from each point in the trajectory. We quantified the endpoint error in a manner agnostic to the exact source of the error (Fig. S7). We looked for any correction that occurred as the hand was decelerated to rest at the end of the reaching movement. Specifically, we calculated the magnitude and sign of the largest lateral deviation from the final hand position, after the hand had exceeded 80% of the target-to-target displacement. This endpoint error could either be positive or negative; our convention is that positive errors indicate errors in the direction of the holding force (should reduce holding force magnitude) and negative errors indicate errors opposite the direction of the holding force (should stabilize and increase the holding force magnitude).

### Familiarization

All human participants were given familiarization trials at the start of each experiment. During this familiarization period, participants learned the desired movement speed through audiovisual feedback and scoring, as described in *Trial Structure*. The experimenter clarified the meaning of all visual stimuli in the workspace and instructed participants to obtain as many points as possible. After participants were able to reach between targets comfortably and accurately, the experiment score was reset and the experimental paradigm began.

After practicing reaching movements, further familiarization was provided to participants who were to perform the working memory task during holding (Experiments 5, 10, and 11). First, these individuals were exposed to the working memory task in isolation (*i.e*., without having to perform reaching movements). Then, a practice period was given where postural probe trials were interspersed within normal reaching trials. For the first few postural probe trials in this practice period, the experimenter warned the participant that the reaching movement would end with the working memory task.

### Neural integration in normal reaching movements at the start of each experiment

In the main text, we start by asking if the integral of the motor commands used to move the arm predict the motor commands that hold the arm still at the end of movement. To answer this question, we considered all trials in the null field periods (the period before the introduction of the force field) of each experiment. In addition to the null field periods within the 15 experiments described below, we included null period trials from an additional experiment (n=23) not described or included in our primary analysis. For this additional experiment, we considered only the relevant initial period which consisted of null field trials. This period consisted of 40 total reaching movements, 20 outwards movements and 20 backwards movements. Every 2^nd^ outwards reach was performed in a channel trial where forces were measured. All backwards trials were performed in channels. The amplitude of each reaching movement was 10 cm, and reaches were performed away from and towards the body.

### Experiments 1-4: Measuring the causal effect of moving on holding

We conducted different experiments to test if the relationship between holding force and the moving force integral was maintained despite changes in the position of the arm in the workspace, the direction of the reaching movement, and the direction of the force perturbation.

In Experiment 1, we tested participants (n=15) in a paradigm consisting of 10 cm reaching movements, where the hand was centered in front of the body and outwards reaching movements were performed directly away from and the body. In Experiment 2, we tested participants (n=11) in a paradigm consisting of 10 cm reaching movements, where the starting position was placed in front of the body but outwards reaching movements were performed at an oblique angle of 135° (away from and to the left of the midline of the body). In Experiments 3 and 4, we tested participants (n=9 for Experiment 3, n=8 for Experiment 4) in a paradigm consisting of 10 cm reaching movements, where the hand was placed to the left and the right of the midline of the body by approximately 15 cm and 5 cm respectively, and outwards reaches were performed only along the y-axis of the workspace away from the body (essentially these experiments were analogues of Experiment 4, translated to the left and right). Experiments 3 and 4 were identical, except that in Experiment 3 participants were adapted to CCW force fields, and in Experiment 4 participants were adapted to CW force fields.

Experiments 1-4 followed the same general trial structure (Fig. 3B). Before exposure to the perturbation, participants reached for 40 trials (20 outwards trials and 20 backwards trials) in a baseline period. Every 2^nd^ outwards reach was performed in a channel where moving and holding forces were measured. Next, the adaptation period started, which consisted of two blocks with 3 phases each. In the first phase, a CW/CCW velocity-dependent perturbation was introduced, and gradually increased/decreased from 0 to 15/-15 N-s/m over the course of 100 outwards reaching trials (200 trials in total). The perturbation magnitude was then maintained at a constant level of 15/-15 N-s/m over the course of 50 outwards reaches (100 trials total) and then brought back to zero gradually in a de-adaptation period of 100 outwards reaching trials (200 trials total). After this de-adaptation period, participants continued to reach in a washout period of 20 outwards reaches (40 trials total) where no force field was applied. Participants were then given a short break and this structure was repeated. In Experiments 1 and 2, participants were exposed to either a CCW or CW perturbation in the 1^st^ block, and the opposite perturbation in the 2^nd^ block (order counterbalanced across subjects). In Experiments 3 and 4, participants were exposed to either the left-most target or the right-most target in the 1^st^ block, and the opposite target in the 2^nd^ block (order counterbalanced across subjects). During the adaptation and de-adaptation periods, we measured subject moving and holding forces in channel trials on every 5^th^ outwards reaching movement.

### Experiment 5: Measuring the duration of holding forces

As participants adapted to a velocity-dependent force field, they also applied large forces as they held their hand still within the target at the end of the reaching movement. We recruited a set of subjects (n=7, Fig. S1) to determine if this holding force was stable in time. We gradually adapted participants to a CCW velocity-dependent force field. At various points during the experiment, we measured holding forces for a long period of time (6.5 seconds in total). To keep participants engaged in the holding process, during the final 6 seconds of the holding period, subjects were distracted with the working memory task (same task used in measuring the postural field). During this period of time an endpoint clamp was engaged to hold the hand at its final position. Endpoint clamps were applied to the hand on all outwards reaching movements, but only lasted 1.8-1.9 seconds on normal reaching trials with no working memory task. The reaching movement preceding the measurement of holding force was performed in a channel. Outwards reaching movements of 10 cm were all performed directly away from the body.

Before adaptation, holding forces were measured in a block consisting of 100 trials (50 outwards reaches and 50 backwards reaches). On every 5^th^ outwards reaching trial, the holding period was extended and the working memory task was presented. After this period, participants took a short break. After the break, the velocity-dependent perturbation was introduced. The field magnitude was decreased from 0 to −7.5 N-s/m in constant increments over the course of 50 outwards reaching trials (100 trials total). Over the next 50 outwards reaches (100 trials total) holding forces were again measured during an extended period on every 5^th^ reach. On all other reaches, the force field was maintained at a constant level (−7.5 N-s/m). Next the force field was gradually deceased further over 50 outwards reaches (100 trials total) to −15 N-s/m. During the final 50 outwards reaches (100 trials total), holding forces were again measured during the working memory task on every 5^th^ reach. On all other reaches, the force field was maintained at a constant level (−15 N-s/m).

### Experiment 6: Reaching from different start locations to the same endpoint

We tested whether different holding forces could co-exist at the same location in space, if the moving forces that brought the hand to that location differed. We first tested this idea by exploiting the phenomenon of spatial generalization^42,43^ where adaptation in one location in space, can generalize to other locations in space where perturbations were never experienced. We designed an experiment, where two starting locations would lead to the same target location, but each reach would generalize moving forces with opposing directions.

The generalization paradigm had one start position, but two target positions. Target 1 was located 10 cm directly away from the body, relative to the start position. Target 2 was located 10 cm directly towards the body, relative to the start position. Participants (n=13) reached in a repetitive 4 trial structure (Fig. S2A): start position to Target 1, Target 1 back to start position, start position to Target 2, and finally Target 2 back to start position. Because of this geometry, generalization of learning could occur because reaches from the start position to Target 1 were made in the same direction as reaches from Target 2 to the start position, and reaches from the start position to Target 2 were made in the same direction as reaches from Target 1 to the start position. We perturbed reaching movements from the start position to Target 1 and Target 2, but never perturbed reaching movements made towards the start position. Instead, these reaches were always made in channel trials.

The experiment paradigm consisted of gradual adaptation to a CW velocity-dependent force field. In other words, reaches to Target 1 were pushed to the right, and reaches to Target 2 were pushed to the left. This presented an interesting scenario (Fig. S2D and S2E), where the forces generalized when reaching to the start position would sometimes be oriented to the right (Target 2 to start), and other times be oriented to the left (Target 1 to start). Participants were adapted to the force field over the course of 100 cycles. In each cycle, all 4 possible reaching movements were completed one time. On every 5^th^ cycle, reaches to Target 1 and Target 2 were made in a channel in order to measure moving and holding forces. The CW perturbation gradually increased from 0 to 15 N-s/m in even increments across trials.

### Experiment 7: Reaching to a single endpoint with variable reaching forces

In Experiment 7, we considered a paradigm in which reach forces differed, but the start and endpoint of the movement was the same. In this experiment, participants (n=14) participated in a dual adaptation paradigm^44^. In this paradigm, rather than controlling a single cursor, participants controlled the motion of a rectangular box centered on the hand (Fig. S6A). On some trials, a control point and virtual target were placed on the left side of the tool. On other trials, a control and virtual target were placed on the right side of the tool. Participants were instructed to move the displayed control point to the displayed target. Critically, even though a different control point was used on alternating trials, the reaching movement started and began at the same locations in space. The control points were instead used as an implicit cue for the direction of an upcoming perturbation. During the adaptation period of the experiment, on trials where Control point 1 was displayed, a CW perturbation was applied to the reaching movement. On trials where Control point 2 was displayed, a CCW perturbation was applied to the reaching movement.

At the start of the experiment, participants reached with each control point a total of 20 times (40 reaches total). Then the CW or CCW perturbations were abruptly increased to 15 or −15 N-s/m, respectively. Participants adapted to the perturbation over the course of 100 Control point 1 reaches and 100 Control point 2 reaches (400 trials total). On every 5^th^ outwards reaching movement, the reach was performed in a channel, and moving and holding forces for either Control point 1 or Control point 2 were measured. Reaching movements were always 10 cm in magnitude, and directed away from or towards the body.

### Experiments 8 and 9: Reaching in a zero integral force field

We considered an alternative hypothesis to neural integration that could explain the causal relationship we observed between moving forces and holding forces; holding forces could be a trivial continuation of the forces produced at the end of movement, just prior to the start of holding. To test this hypothesis, we designed a position-dependent force field, composed of two components (Eqs. 1.2 and 1.3): one that acted on the first half of the reaching movement (FF_1_, first 10 cm of the reach) and one that acted on the second (FF_2_, last 10 cm of the reach). By adapting individuals to a force field composed of both FF_1_ and FF_2_, we created a scenario where large forces were produced at the end of the reaching movement, but the reaching forces integrated to zero.

Throughout Experiment 8 (n=14 subjects) we measured moving and holding forces in channel trials on every fifth outwards reaching movement. The experiment started with 25 unperturbed outwards reaching trials (50 trials total). After this, we gradually adapted participants to a force field that only perturbed the hand during the second half of the reaching movement (Figs. 4A and 4B). To do this, we gradually increased the magnitude of FF_2_ from 0 N to 3.5 N in even increments, over the course of 100 outwards reaching movements (200 trials total). During this time, the magnitude of FF_1_ was maintained at 0 N. We reinforced this learning period with an additional 10 outwards reaches (20 trials total) where FF_2_ was maintained at 3.5 N and FF_1_ at 0 N. In the final part of the experiment, we maintained FF_2_ at 3.5 N, but gradually decreased FF_1_ from 0 N to −3.5 N, to teach individuals a new set of forces during the first half of the reaching movement. We adapted individuals to this bidirectional force field slowly by decreasing FF_1_ to its terminal magnitude over the course of 200 outwards reaching trials (400 trials total). In this way, at the end of the experiment, participants were exposed to two force fields within the same reaching movement that perturbed the hand in opposite directions but with equal magnitude.

We found that holding forces gradually decreased in the final phase of the experiment as individuals adapted to FF_1_, consistent with our hypothesis of integration. If adaptation to FF_1_ was the sole driver of the reduction of holding force, if FF_1_ was omitted entirely, the holding forces should not decrease. To test this idea (and also confirm that the reduction in holding force was not caused by others factors like fatigue or accumulation of endpoint errors), we recruited a control group (n=11). The control experiment (Experiment 9) also started with 25 unperturbed outwards reaching trials (50 trial total). As in Experiment 8, control participants were then adapted to FF_2_ only. The magnitude of FF_2_ was gradually increased from 0 to 3.5 N over the course of 100 outwards reaches (200 trials total), while FF_1_ was maintained at 0 N. During the final phase of the experiment, FF_2_ was maintained at 3.5 N and FF_1_ was maintained at 0 N for an additional 210 outwards reaches (420 trials in total, to match the trial count of Experiment 8).

### Experiment 10: Altering the reach forces and measuring its effects on the postural field

Experiments 1-9 suggested that holding forces were causally coupled to moving forces, but did not explain how holding forces contribute to the maintenance of a stable arm posture. To determine the relationship between holding forces and the final arm posture, we systematically displaced the hand during the holding period in a random direction (see *Measuring the postural field*). We used these probes to measure the restoring force participants applied to the handle in response to the unintended displacement. These forces formed a set of measurements that constituted a postural field (Fig. 5B). We measured the postural field before and after adaptation to a velocity-dependent force field.

To determine if the postural field changed after changes to the reaching forces, we adapted a set of participants (n=27) to a CCW velocity-dependent force field. To measure the postural field in two-dimensional space, postural probes moved the hand in 1 of 12 directions (0°, 15°, 30°, 45°, 90°, 135°, 180°, 225°, 270°, 315°, 330°, and 345° with respect to the x-axis, while participants were distracted with a working memory task (see *Working Memory Task*). We measured the postural field before and after adaptation. Before adaptation, participants completed 3 blocks of trials, each separated by a short break. In each block, all 12 postural probe directions were visited a single time. The probe displacement was 2.5 cm for all probe directions. Postural probes were given on every 4^th^ outward reaching movement. Therefore, participants completed a total of 288 baseline trials (3 blocks x 12 probes/block x 4 reaching movement pairs/probe x 2 movements/reaching movement pair). Outwards reaching movements of 10 cm were all performed directly away from the body.

Participants were then gradually adapted to a velocity-dependent force field. The field magnitude (*f* in Eq. 1.1) was decreased from 0 to −10 N-s/m in constant increments over the course of 65 outwards reaching trials (130 trials total). After this adaptation period, the postural field was re-measured. As before, all 12 probe directions were probed in a random order, 3 times. No breaks were provided in between blocks. We anticipated that the postural field would shift after adaptation to the force field. Therefore, we extended the probe displacement to 4 cm for probe angles of 0°, 15°, 30°, 45°, 315°, 330°, and 345°. Postural probes occurred at the same frequency as before adaptation for a total of 288 trials. To maintain participants in an adapted state, on all outwards reaching trials other than postural probe trials, a velocity-dependent perturbation was maintained at −10 N-s/m. Endpoint clamps were applied to the final hand position during all outwards trials (with the exception of postural probe trials) during the holding period (1.5 second duration, with an addition 0.3-0.4 inter-trial-interval).

To quantify the postural field before and after adaptation (Fig. 5E), we fit a two-dimensional linear spring (Eq. 4) to x and y forces measured on postural probe trials for each subject individually. The model and fitting process are described in *Quantifying the null point and shape of the postural field*. For visualization purposes (Fig. 5B), we constructed a two-dimensional postural field from the forces measured during probe trials using linear interpolation. To do this, along each probe direction we resampled forces in x and y spatially in intervals of approximately 0.1 cm. For each of the resampled restoring forces we calculated the corresponding polar coordinates (*i.e*., the radius and angle). In polar coordinates, all x and y forces lied along a rectangular grid. We used bilinear interpolation along these polar coordinates to estimate the postural field within the space between the 12 probe angles.

### Experiment 11: Measuring the relationship between holding forces and the null point of the postural field

We reasoned that if holding forces truly reflected misalignment between the position of the hand and the null point of the postural field, we could gradually eliminate these forces if we displaced the hand towards its null point. We recruited a set of subjects (n=19) to test these predictions throughout the process of gradual adaptation to a velocity-dependent force field. Given our findings in Experiment 10 we expected that the null point of the postural system would be located laterally to the hand in the direction of the compensatory hand force. For this reason, to measure forces in the direction of the null point, we exposed participants to probes along this single direction (0° with respect to the x axis). As always, during the postural probe, participants were distracted with the working memory task. For the first ten participants, we used 5 cm postural probe trials. For the last nine participants, we shortened the probe length to 4 cm. Here we analyze only the first 4 cm of displacement to combine both versions of the experiment.

Before adaptation, we measured the postural forces a total of 10 times. Postural probes were inserted regularly on every 5^th^ outward reach, for a total of 100 trials in this baseline period (10 probes x 5 outwards reaches/probe x 2 reaches/outward reach). Outwards reaching movements of 10 cm were all performed directly away from the body. Next, we adapted participants gradually to a CCW velocity-dependent force field, where we decreased the field magnitude (*f* in Eq. 1.1) from 0 to −10.5 N-s/m in constant increments over the course of 175 outwards reaching trials (350 trials total). We measured hand forces evoked by the postural probe on every 5^th^ outwards reaching movement. On postural probe trials, reaches were performed in a channel.

We found that hand forces during the postural probe resembled a linear spring throughout the process of adaptation. To determine the location of the null point on a trial-to-trial basis we fit a line to the hand forces as a function of hand displacement in the probe, and recorded the x-intercept of the line. To do this, we first resampled subject forces spatially in increments of 0.05 cm. Next, to reduce noise inherent in the single trial force measurements, we used a bootstrapping approach. On each trial, we randomly sampled subjects with replacement, calculated the mean postural force as a function of distance across the sample, and fit a line to this mean behavior. We repeated this process 2000 times, and used this distribution to estimate 95% confidence intervals around the mean (Fig. 5H).

### Experiment 12: Holding forces disappear without endpoint clamps

Our findings in Experiments 10 and 11 presented a paradox. Adaptation of the reach controller changes reaching forces that are integrated downstream into a postural null position that no longer coincides with the desired target. In other words, integration of adapted reaching commands no longer specifies the correct holding location. To solve this potential holding instability, the postural controller must also be adaptive. In all of our experiments (with the exception of Experiment 12) we tried to prevent postural adaptation by clamping the endpoint of each reaching movement with a two-dimensional endpoint clamp. Removal of the endpoint clamp should promote larger endpoint errors, which in turn will cause adaptation in postural control. If correct, this adaptation will bring about an uncoupling between the moving forces and holding forces; as the moving forces increase, the postural controller should learn to no longer apply holding forces.

In search of evidence for this process of adaptation, we recruited a set of participants (n=20) that adapted to a velocity-dependent force field, without any endpoint clamps. The experiment paradigm consisted of two reaching blocks with a short break in the middle. In the first block, before exposure to the perturbation, participants reached for 40 trials (20 outwards trials and 20 backwards trials). Every 2^nd^ outwards reach was performed in a channel where moving and holding forces were measured. Next, we perturbed participants with a 15 or −15 N-s/m velocity-dependent perturbation. To promote larger endpoint errors, this perturbation was introduced abruptly, as opposed to gradually. Participants adapted to this constant perturbation for 200 outwards reaching trials (400 trials total). At the end of the block, the perturbation was abruptly removed, and subjects reached in the absence of the force field for an additional 45 outwards trials (90 trials total). The second block followed the same structure as the first block. The blocks differed in the direction of the force field (CW or CCW) which was counterbalanced across participants.

### *Experiments 13* and *14: Large endpoint errors lead to a reduction in holding forces*

Experiment 12 demonstrated that endpoint errors promoted adaptation in the postural controller. In search of additional evidence for postural adaptation, we recruited two sets of subjects (n=15 for Experiment 13 and n=17 for Experiment 14). We gradually adapted participants to a velocity-dependent force field (CCW field for Experiment 13, CW field for Experiment 14). Then, we maintained the perturbation at a constant level for a long period of time. Reaches of 10 cm were performed at oblique angles of 135° as in Experiment 2. Along this reaching direction, CCW and CW perturbations led to large differences in endpoint errors (Fig. S7A-C). We wanted to know if these different endpoint errors resulted in changes in the size of holding forces, reflecting the process of adaptation.

Experiments 13 and 14 both followed the same trial structure. Before exposure to the perturbation, participants reached for 40 trials (20 outwards trials and 20 backwards trials) in a baseline period. Every 2^nd^ outwards reach was performed in a channel where moving and holding forces were measured. Next, the adaptation period started, which consisted of 2 phases. In the first phase, a CW/CCW velocity-dependent perturbation was introduced, and gradually increased/decreased from 0 to 15/-15 N-s/m over the course of 100 outwards reaching trials (200 trials in total). In the second phase, the perturbation magnitude was then maintained at a constant level of 15/-15 N-s/m over the course of 200 outwards reaches (400 trials total). We measured subject moving and holding forces in channel trials on every 5^th^ outwards reaching movement.

### Experiment 15: Measuring cortical involvement in the control of reaching and holding

Altogether, our findings indicated that the control of reaching was dependent on the process of moving; forces that moved the arm were integrated into a null point that was then applied to postural control at the end of movement. Like the eye, our experiments suggested that the moving controller and the holding controller were connected in serial. For the oculomotor system, move and hold commands are generated by separate regions in the brainstem. We wondered if this is also true in the control of reaching and holding.

We recruited patients (n=13) who had survived a stroke affecting cortical or subcortical white matter associated with the corticospinal tract (CST). We measured the degree of motor impairment in the patients using the Fugl-Meyer Assessment (FMA) and the Action Research Arm Task (ARAT). Two separate raters scored each assessment and scores were averaged across raters. In each limb, we measured the strength of elbow flexion/extension and shoulder horizontal adduction/abduction using a dynamometer (microFet 2). During measurements, participants rested their arm on a side table which supported their arm so it rested slightly below shoulder level. Strength measurements were repeated twice, the maximal force was recorded on each effort, and forces were averaged over repetitions.

During each measurement, patients were verbally encouraged by the experimenter to produce as much force as possible. FMA scores, ARAT scores, strength, and other patient characteristics are reported in Table S1. Missing entries in table indicate that the patient was unable to perform the desired action. Patients were selected based on MRI or CT scans, and/or available radiologic reports. Scans and/or reports were corroborated to determine the level at which the white matter of the corticospinal tract (CST) was lesioned. Table S1 provides the level of the brain at which the white matter was lesioned.

If moving used cortical control, and the holding integrator used subcortical control, we reasoned that stroke patients may show impairment in the generation of reaching movements, but the process of integrating moving forces into holding commands would be spared. We tested the stroke patients and a set of healthy age-matched controls (n=10) on a paradigm similar to Experiment 1. To account for motor impairment, we relaxed the timing requirements of our standard reaching experiment. Participants were awarded a point for reaches (10 cm in magnitude) that terminated within 600-800 ms (as opposed to 450-550 ms). Additionally, the arm of the patient was supported against gravity by an air sled that rested on a smooth table located underneath the visual display. Age-matched controls were tested in an identical protocol.

Each experiment block started with 20 unperturbed outwards reaches (40 trials total). After this initial period, we gradually introduced a CW or a CCW velocity-dependent force field. Over the course of 80 outwards reaches (160 trials total) we gradually increased (CW) or decreased (CCW) the force field magnitude in even increments from 0 N-s/m to 18 or −18 N-s/m. The force field magnitude was maintained for an additional 40 outwards reaches (80 trials total) and then brought gradually back to 0 N-s/m over the course of 80 outwards reaches (160 trials total). The experiment block ended with 20 unperturbed outwards reaching movements (40 trials total). We measured moving and holding forces on channel trials on every 5^th^ outwards reaching movement.

We measured behavior in both the paretic (contralateral to the lesion) and non-paretic arms. In this way, we could also determine the effect of paresis on reaching and holding forces within each patient. The paretic and non-paretic arms were tested in different blocks, with 2 blocks for each arm.

The order of the blocks was always paretic, then non-paretic, then paretic, then non-paretic. In each block either a CCW or CW force field was applied. Across blocks, the perturbation order was always A-B-B-A where A and B refer to either a CW or CCW force field. The force field orientation on the first block was counterbalanced across participants. Healthy age-matched controls were tested in a similar manner. In the absence of a paretic/non-paretic distinction, we tested either the left or the right arm in the first block (counterbalanced across participants).

To identify any impairment in reaching we focused on various kinematic measures. In all of these measures, we only considered trials during the null field period prior to the introduction of the force field. First, we considered time-courses for how far the arm deviated from a straight-line trajectory. For this, we calculated the unsigned lateral deviation from the straight-line trajectory for each trial in the null field period, averaged these time-courses across trials within each subject, and then finally averaged these mean traces across subjects (Fig. 6C). In addition to this, we also calculated several kinematic measures of performance: the duration of the reaching movement (Fig. 6D) and the displacement from the target at reach endpoint (Fig. 6E).

We found that erratic reaching forces on null field trials led to sustained holding forces (Fig. 6H). To determine if coupling between moving and holding forces is affected by stroke, we selected the most impaired reaching movements. To do this, we identified all reaches where the integral of moving forces fell outside of 2 standard deviations of the moving force time-integral distribution of the healthy control population (Fig. 6G). We analyzed if these highly impaired reaching forces were integrated similarly to normal reaching movements (Fig. 6H). To do this, we separated trials into bins according to the integral of the moving force.

To further search for an impairment in the process of neural integration, we looked at the gain of the linear relationship between reaching and holding, after exposing participants to a velocity-dependent curl field. We fit a linear model to the trial-by-trial measurements of holding forces and moving force integrals for each individual participant (Fig. 6J) and tested if this gain differed across the paretic arm of patients, non-paretic arm of patients, left arm of healthy control participants, and right arm of healthy control participants (Fig. 6L). When collapsing moving and holding forces across participants (*e.g*., Fig. 6K) we accounted for the increased movement variability observed in the paretic arm by first sorting the trial-by-trial behavior with respect to the magnitude of the moving force integral, before averaging moving and holding forces across participants (within 1 trial bins).

---

**Figure S1.**
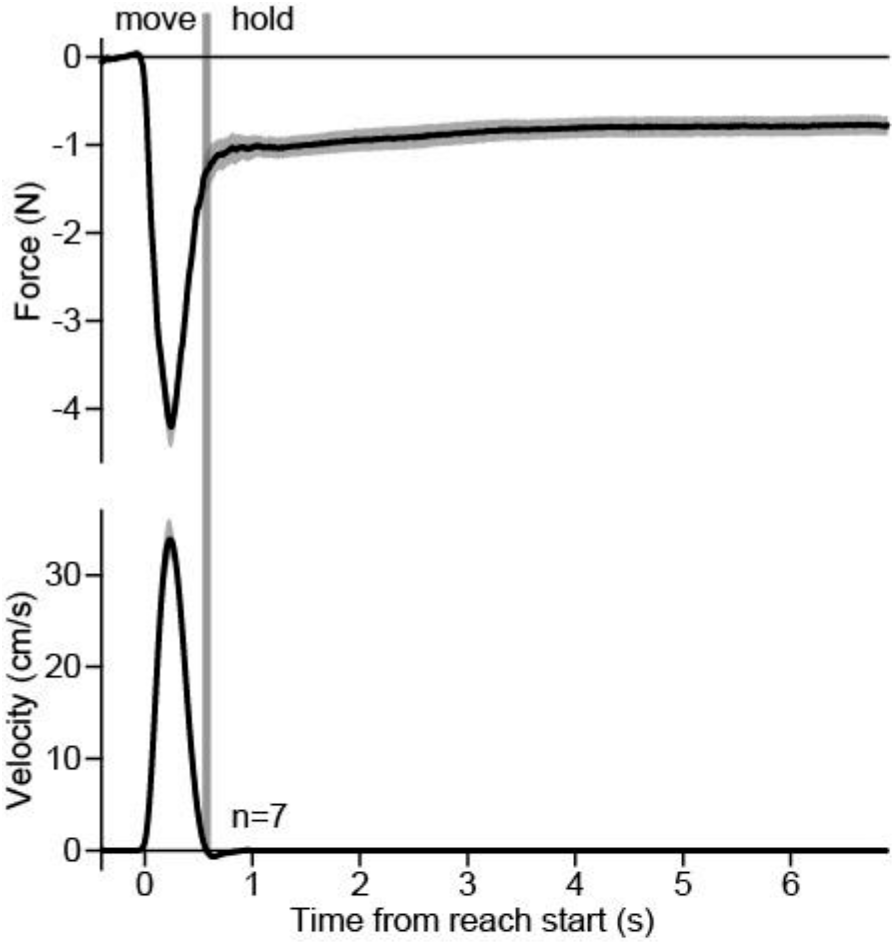
Holding forces are sustained across long time intervals. Participants (n=7) were exposed to a CCW velocity-dependent curl force that increased in magnitude across many trials. Here we show the lateral force (top) and hand velocity (bottom) after exposure to maximal field strength. The vertical line denotes the end of the movement, hence, the start of the holding period. On the trials shown, participants held the hand at the target and waited for a long inter-trial-interval to start the next reaching movement. During this period of time, participants engaged in a working memory task.

**Figure S2.**
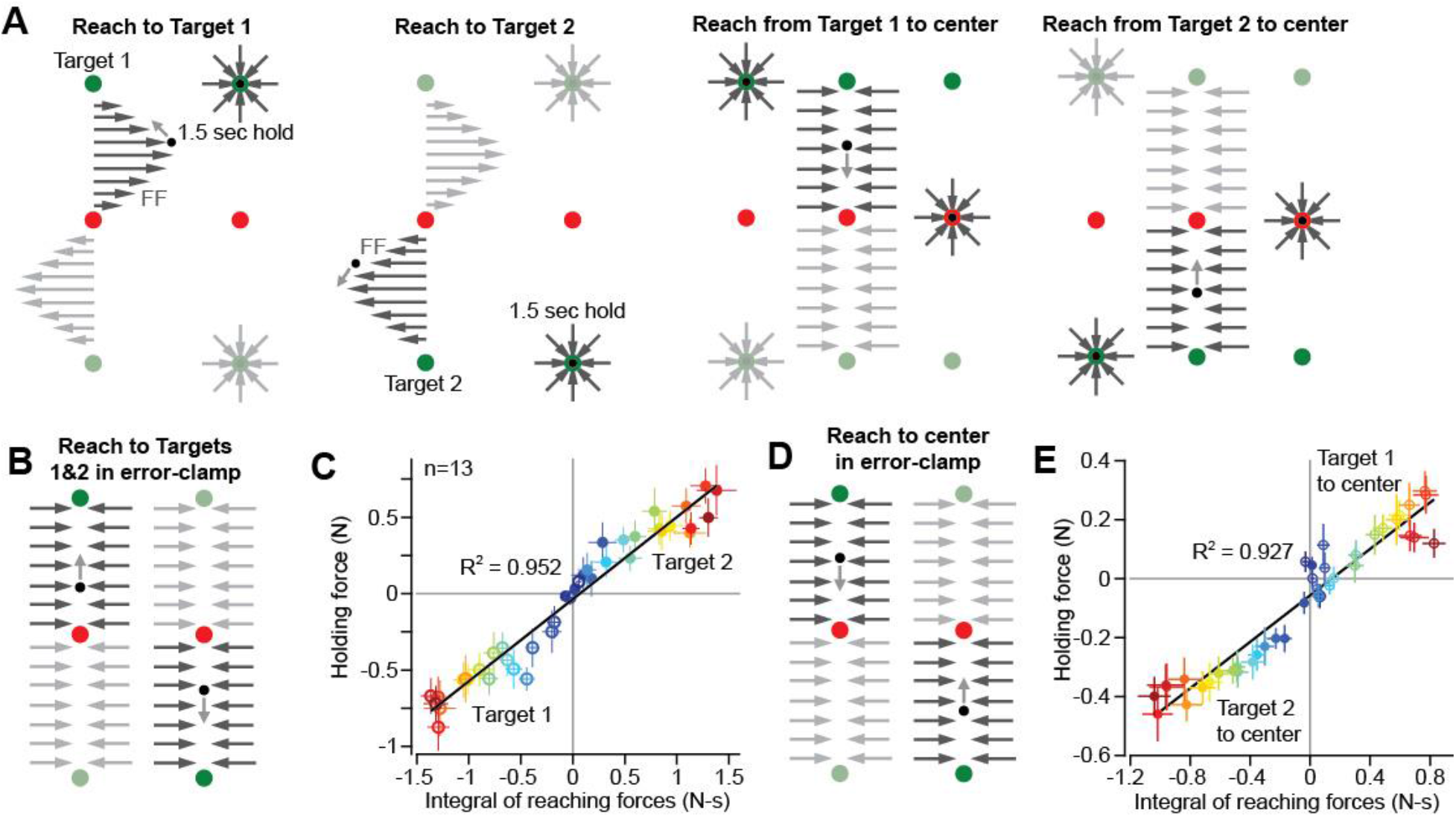
Holding forces at the same target position are specific to reaches initiated from different start positions. **A.** Participants (n=13) reached in cycles of four trials. On the first trial (first column), participants reached away from the body from a center position (red) to Target 1 (green). At the end of the movement, the hand was held in place for a long holding period. On the next trial (third column), the participant returned the hand from Target 1 to center. On the third trial of the cycle (second column), participants reached from the center position to Target 2. And finally, on the fourth trial (fourth column), participants returned the hand back to center again. After an initial familiarization period, participants were exposed to force fields when reaching from the center to Target 1 (rightwards field) and from the center to Target 2 (leftwards field). Reaching movements that terminated at the center position were always executed in a channel that prevented lateral errors. **B.** Occasionally, channel probe trials were inserted to measure hand forces when reaching to Targets 1 and 2. **C.** Shown is a scatterplot of the holding force as a function of the moving force time-integral for the probe trials depicted in B. Each point represents one trial. Color indicates the strength of the force field at that point in the experiment (blue means small, and red means large). Unfilled circles show reaches to Target 1 and filled circles show reaches to Target 2. The solid line shows the linear regression across trials. Values are mean ± SEM across participants. **D.** All reaching movements made towards the center position were executed in a channel. **E.** Same as in **C** but for probe trials depicted in **D.** Color indicates the force field magnitude at a specific point in the experiment. Filled circles depict reaching movements from Target 2 to center and unfilled circles depict reaching movements from Target 1 to center. Note that this implies that different holding forces coexisted at the same location in space (the center) on different trials, but at the same point in the experiment. The solid line shows the linear regression across trials. Values are mean ± SEM across participants.

**Figure S3.**
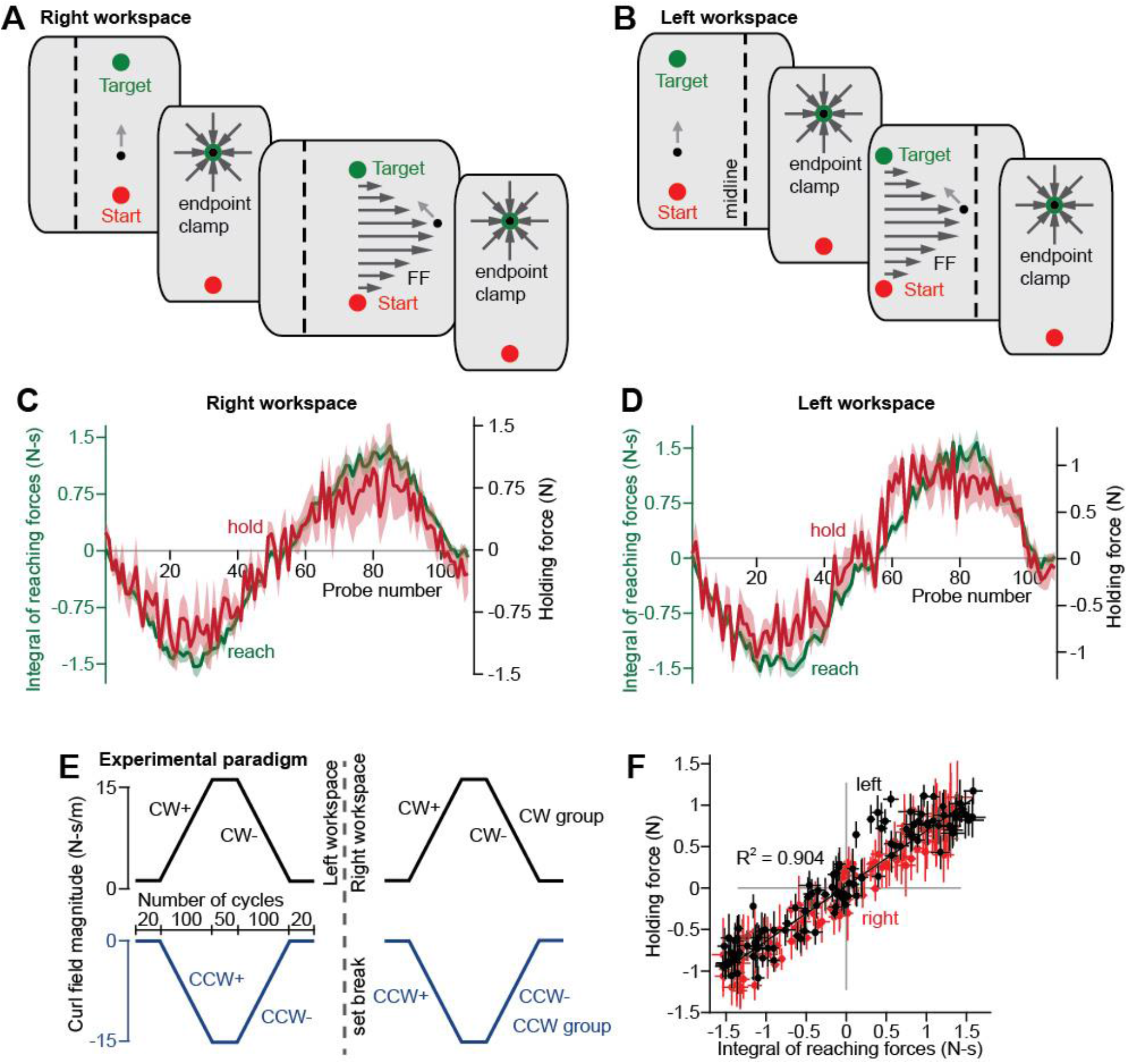
The relationship between reaching and holding forces is consistent across different arm postures. **A.** The movement was centered about 15 cm to the left of the midline of the body (dashed line). First, participants reached in a null field (top) but were then exposed to a CCW (Experiment 3, n=9) or CW (Experiment 4, n=8) force field that gradually increased in strength over many trials. At the end of each null field or perturbation trial, the hand was held in place during an extended holding period within an endpoint clamp. **B.** Participants also reached in a separate block in the same manner as in **A**, but with the arm centered 5 cm to the right of the midline of the body. **C.** Occasional probe trials were inserted to measure hand forces in the left workspace. The time-integral of the reach force is shown on each trial in green. The holding force is shown in red. The + and - indicators represent periods of time in which the force field was increasing and decreasing, respectively. Values are mean ± SEM across participants. **D.** Same as in C but for the right workspace. **E.** Participants were gradually adapted and then deadapted to either a CCW (blue group, bottom) or CW (black group, top) force field. Participants were tested in two blocks separated by a short break. In one block, participants reached with the arm centered to the left of midline (shown to the left of dashed line). In the other block, participants reached with the arm centered to the right of midline (shown to the right of dashed line). The order was counterbalanced across participants. **F.** Shown is a scatterplot plot depicting the holding forces as a function of the moving force time-integrals. Points represent individual trials. The black line shows the linear regression. Values are mean ± SEM across participants.

**Figure S4.**
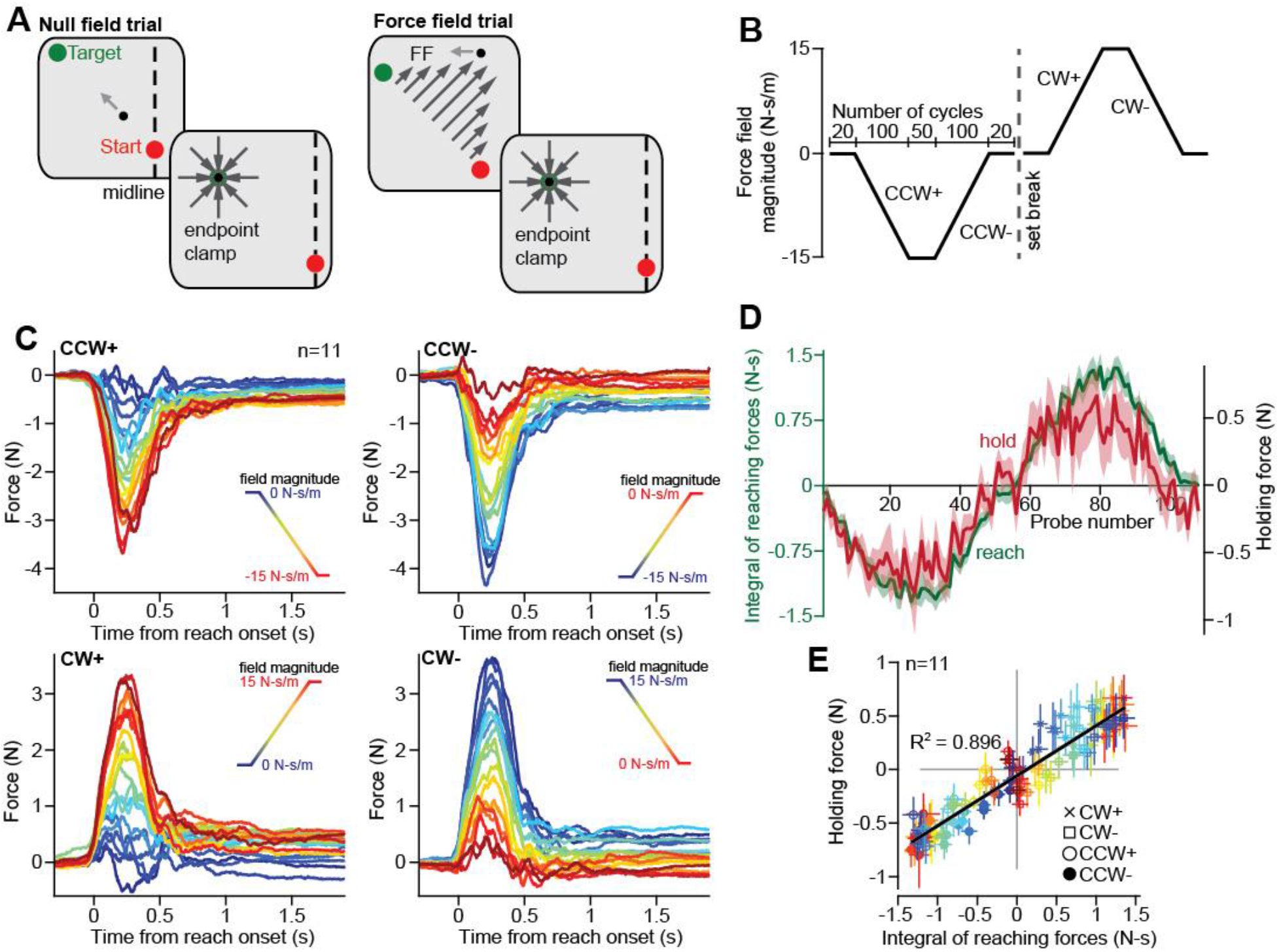
Moving and holding forces for reach movements of different angles. **A.** Participants (n=11, Experiment 2) reached between a start location and a target located at an oblique angle (135° with respect to the positive x-axis) from the midline of the body (represented by the dashed line). First, participants reached in a null field (left), but then were exposed to CW or CCW velocity-dependent force fields (right). At the end of null field and force field trials, the hand was held in place with an endpoint clamp. **B.** The force field schedule. Shown is the force field magnitude on each trial. The experiment was composed of cycles consisting of one outwards reach and backwards reach. On outwards trials were perturbed. The force field gradually increased to a constant value and then decreased. The + and - indicators represent periods of increasing and decreasing force field strength, respectively. CCW and CW force fields were applied in different blocks, separated by a short break. **C.** Occasionally, probe trials were inserted to measure forces during moving and holding. Shown are forces during different periods of the experiment (upper-left is increasing CCW field, upper-right is decreasing CCW field, lower-left is increasing CW field, and lower-right is decreasing CW field). Each trace is one trial. The color indicates the field magnitude in each respective period (blue is early and red is late). **D.** We calculated the time-integral of the moving forces (green) and the static holding force (red) on each trial. Values are mean ± SEM across participants. **E.** Shown is a scatterplot of the holding forces as a function of the moving force time-integrals. Colors are consistent with trials in **C.** Marker shape depicts the period of the experiment (legend bottom-right). The solid line depicts the linear regression. Values are mean ± SEM across participants.

**Figure S5.**
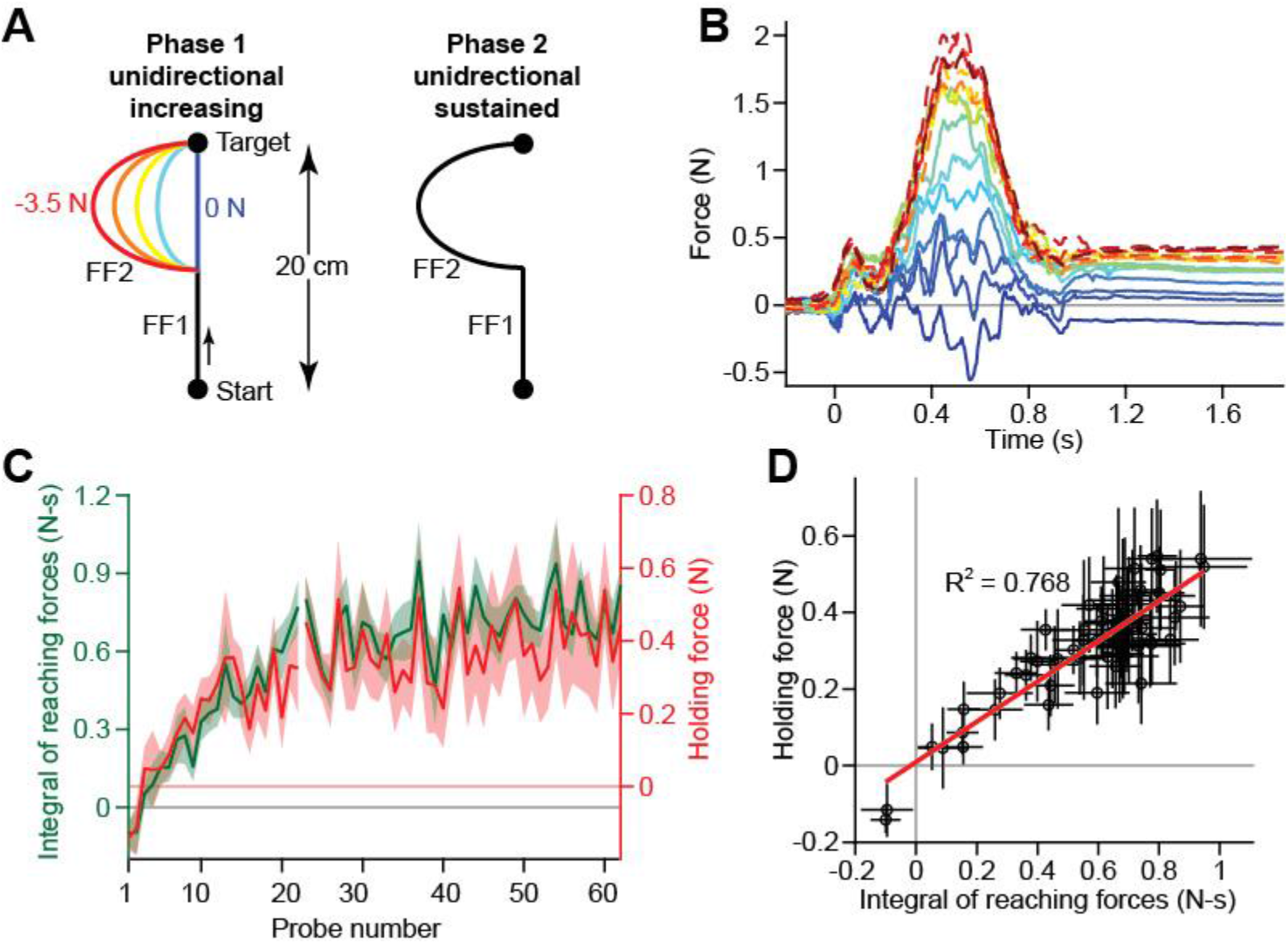
Holding forces in a sustained position-dependent force field. **A.** Participants (n=11, Experiment 9) reached between two locations separated by 20 cm. In the first phase of the experiment, participants were exposed to a position-dependent force field that was only active during the second half of the movement. The magnitude of the force field was increased over several trials (indicated by color, blue to red). In the second phase of the experiment, the perturbation was maintained for many trials. **B.** Occasionally, probe channel trials were inserted to measure hand forces. Shown are hand forces in time over sixteen different trial bins (roughly four probe trials in each bin). Blue to red denotes trials from early to late in the experiment. Solid traces correspond to Phase 1 and dashed traces correspond to Phase 2. **C.** We calculated the time-integral of moving force (green) and the final holding force (red) on each trial. Values are mean ± SEM across participants. **D.** Shown is a scatterplot of the holding forces as a function of the moving force time-integral. Each point is a trial. The solid red line depicts the linear regression. Values are mean ± SEM across participants.

**Figure S6.**
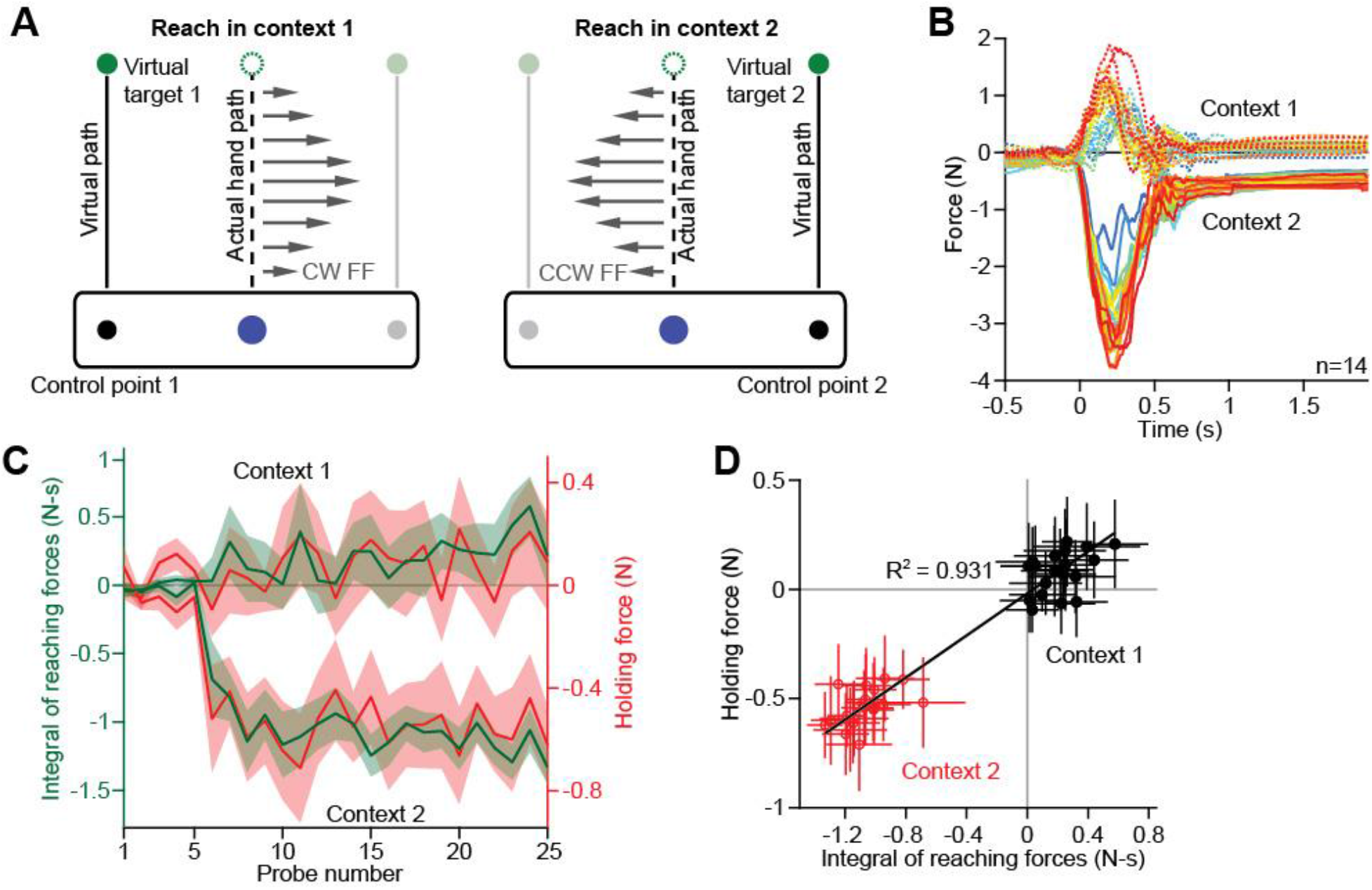
Moving and holding forces during exposure to dual force fields. **A.** Participants (n=14, Experiment 7) reached between two locations in space (the filled blue circle and the dashed central green circle). Rather than controlling a single cursor, participants were shown a rectangular tool that was centered on the hand. On some trials (context 1, left schematic), a control point appeared on the left of the tool and participants were told to move that point into a target that appeared to the left in the workspace. On other trials (context 2, right schematic), a control point appeared on the right of the tool and participants were told to move that point into a target that appeared to the right in the workspace. Critically, the physical starting position and target position were the same for all of the trials, simply the context changed. Participants were exposed to CW and CCW force fields. The orientation of the field was paired to the context (context 1 paired to CW field and context 2 paired to CCW field). **B.** Forces were measured on occasional channel probe trials. Here we show the forces produced on probes in context 1 (dashed lines) and context 2 (solid lines). Color indicates the trial number (blue is early in the experiment, red is late). **C.** We calculate the moving force time-integral (green) and holding force (red) for context 1 trials (top) and context 2 trials (bottom). Values are mean ± SEM across participants. **D.** Next we examined the scatterplot showing the holding force one each trial as a function of the moving force time-integral on that trial. Black points correspond to context 1 trials and red points correspond to context 2 trials. To reiterate, all of the trials start and end at the same location in space. The solid black line shows the linear regression. Values are mean ± SEM across participants.

**Figure S7.**
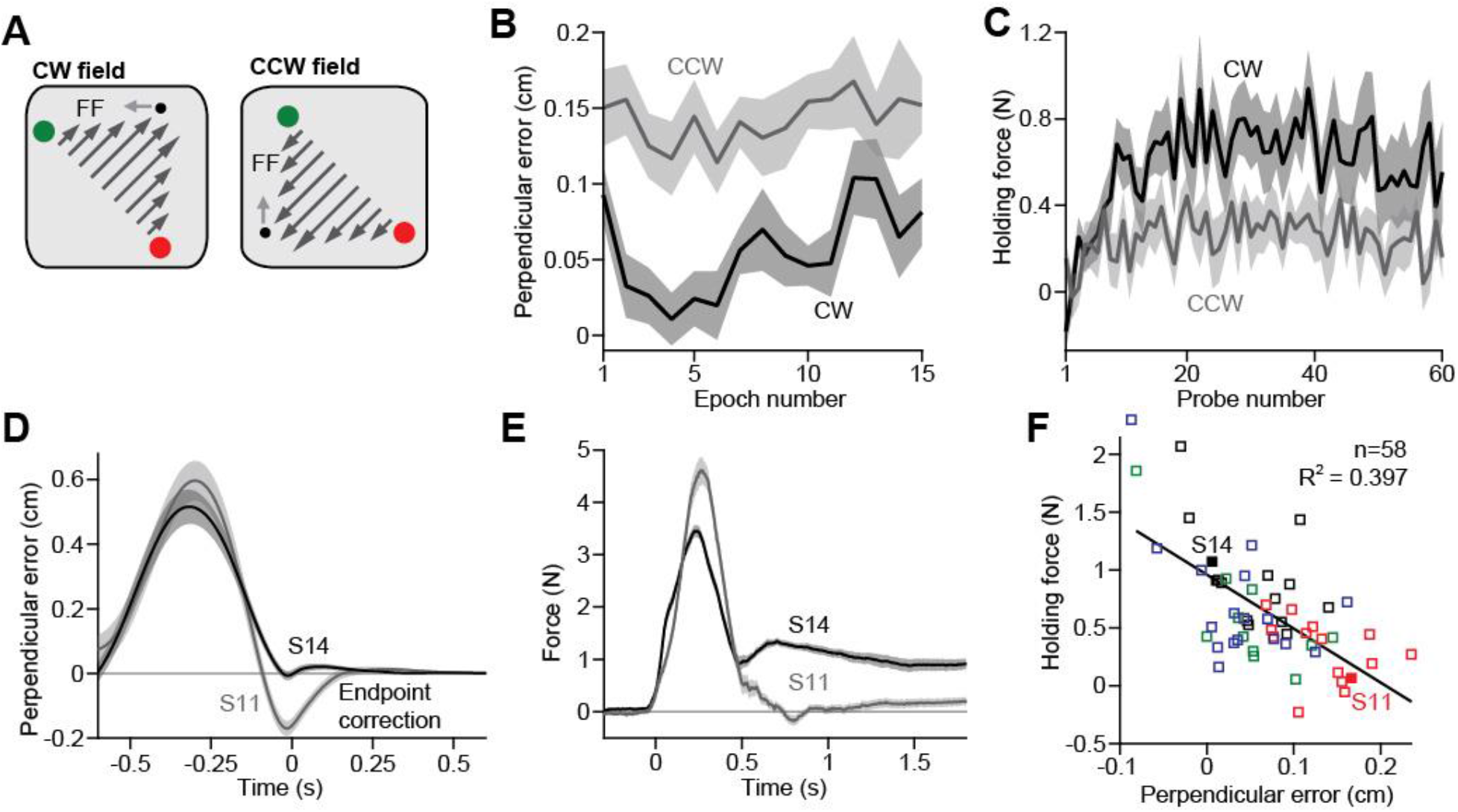
The neural integrator adapts to positional errors. **A.** Participants reached between two oblique targets and were gradually adapted to either a CCW (n=15, Experiment 13) or CW (n=17, Experiment 14) velocity-dependent curl field. **B.** Compensatory forces led participants to make positional errors while attempting to decelerate and stop the hand on the target. To quantify these errors, we measured the maximum deviation of the hand from the straight-line path, in the late part of the movement after 80% of the reaching trajectory had elapsed. Positive values indicate errors in the direction of compensatory forces (opposite the robot forces). Due to the inertial properties of the robot and the arm, late reach termination errors were much larger in the CCW field than CW field. Values are mean ± SEM across participants. Each point represents a bin consisting of 16 outwards movements. **C.** We measured holding forces on channel trials interspersed across the experiment. Larger reach termination errors in the CCW field led to smaller holding forces, in a process resembling error-based learning in the postural controller. Values are mean ± SEM across participants. **D.** Here we show the lateral deviation as for two example participants. Traces were aligned to the maximal holding-related error. Participant S11 experienced a much large error in position while attempting to stop the hand at the target than Participant S14. Values are mean ± SEM across trials. **E.** Same as in **D** but for moving and holding forces. Participant S14 had larger holding forces despite smaller moving forces than S11, likely because the postural system learned to eliminate holding forces due to holding-related errors. Values are mean ± SEM across trials. **F.** We collected participants across four experiments (colors denote different experiments) where perturbations were introduced gradually and at equal rates. The y-axis shows the holding force during peak force perturbations. The x-axis shows the perpendicular error on reaching trials that preceded the peak force perturbations. Positive values indicate errors in the direction of holding forces. Each point represents a different participant. The solid black line shows the linear regression across participants.

**Supplementary Table 1.** Measures of impairment in stroke patients. Patients completed the Fugl-Meyer Assessment (FMA) and the Action Research Arm Test (ARAT), as well as strength testing in the shoulder and elbow. Strength measurements were repeated twice and the maximal force was recorded on each effort and then averaged across repetitions. Two separate raters scored the FMA and ARAT assessments, and scores were averaged across raters. Missing entries in table indicate that the patient was unable to perform the desired action. Patients were selected based on MRI or CT scans, and/or available radiologic reports. Scans and/or reports were corroborated to determine the level at which the white matter of the corticospinal tract (CST) was lesioned. Here we provide the level of the brain at which the white matter was lesioned.

| ID | Age | Sex | Time since stroke | Handed -ness | Paretic arm | Elbow strength (N) |  | Shoulder strength (N) |  | FMA (/66) | ARAT (/57) | CST lesion location |
| --- | --- | --- | --- | --- | --- | --- | --- | --- | --- | --- | --- | --- |
|  |  |  |  |  |  | flexion | extension | adduction | abduction |  |  |  |
| S001 | 80 | M | 2y | Right | Left | 155.2 (P)<br>168.1 (NP) | 82.3 (P)<br>73.0 (NP) | 87.6 (P)<br>103.2 (NP) | 115.7 (P)<br>108.5 (NP) | 57.5 | 57 | Right internal capsule |
| S002 | 51 | M | 6y | Right | Left | 177.5 (P)<br>323.4 (NP) | 114.5 (P)<br>204.6 (NP) | 229.5 (P)<br>345.6 (NP) | 152.6 (P)<br>209.1 (NP) | 40 | 47.5 | Right fronto-parietal white matter |
| S003 | 68 | F | 7y | Right | Right | 142.8 (P)<br>225.5 (NP) | 99.4 (P)<br>153.0 (NP) | 128.1 (P)<br>120.5 (NP) | 107.6 (P)<br>144.1 (NP) | 34.5 | 19 | Left corona radiata |
| S004 | 30 | F | 5y | Right | Left | 116.1 (P)<br>157 (NP) | 83.6 (P)<br>94.3 (NP) | 95.2 (P)<br>119.7 (NP) | 84.5 (P)<br>111.2 (NP) | 55.5 | 43.5 | Right precentral and postcentral gyri |
| S005 | 78 | M | 13mo | Right | Right | n/a | n/a | n/a | n/a | 43.5 | 34 | Left corona radiata |
| S007 | 54 | F | 2mo | Left | Right | 140.6 (P)<br>144.6 (NP) | 101.9 (P)<br>120.1 (NP) | 107.6 (P)<br>109.4 (NP) | 97 (P)<br>85.4 (NP) | 63 | 57 | Left corona radiata |
| S008 | 53 | F | 14mo | Right | Left | 104.5 (P)<br>136.6 (NP) | 65.4 (P)<br>148.1 (NP) | 88.1 (P)<br>133 (NP) | 91.6 (P)<br>125.9 (NP) | 41 | 25 | Right centrum semiovale |
| S010 | 70 | M | 5y | Right | Left | 90.7 (P)<br>197.1 (NP) | 68.1 (P)<br>161.9 (NP) | 177.5 (P)<br>194.4 (NP) | 82.7 (P)<br>144.1 (NP) | 20 | 12 | Posterior limb of right internal capsule |
| S011 | 43 | F | 20mo | Right | Right | 116.5 (P)<br>130.8 (NP) | 97 (P)<br>109 (NP) | 86.7 (P)<br>91.6 (NP) | 98.3 (P)<br>113.9 (NP) | 64 | 57 | Left corona radiata |
| S012 | 48 | M | 6y | Right | Left | 145.9 (P)<br>358.1 (NP) | 91.2 (P)<br>207.3 (NP) | 178.8 (P)<br>292.2 (NP) | 97.4 (P)<br>226 (NP) | 18.5 | 6.5 | Right corona radiata |
| S013 | 68 | M | 9y | Right | Left | 217.1 (P)<br>355 (NP) | 112.5 (P)<br>236.2 (NP) | 220.2 (P)<br>282 (NP) | 130.3 (P)<br>174.4 (NP) | 14 | 8 | Right internal capsule |
| S014 | 45 | F | 16mo | Right | Left | 29.8 (P)<br>114.8 (NP) | 33.4 (P)<br>113 (NP) | 48.9 (P)<br>120.5 (NP) | 46.7 (P)<br>87.2 (NP) | 40 | 39.5 | Right precentral and postcentral gyri |
| S015 | 64 | F | 10y | Right | Left | 149.9 (P)<br>138.78(NP) | 44 (P)<br>140.1 (NP) | 73.4 (P)<br>100.5 (NP) | 48.9 (P)<br>84.1 (NP) | 22 | 4.5 | Right corona radiata |

